# Allosteric regulation of BH3-in-groove interactions by tail anchors of BCL-xL complexes limits BH3 mimetic antagonism

**DOI:** 10.1101/2024.10.02.616265

**Authors:** Laurent Maillet, Aurélie Fétiveau, Lisenn Lalier, Sophie Barillé-Nion, Catherine Guette, Fabien Gautier, Stéphane Téletchéa, Philippe Paul Juin

## Abstract

**In brief:** The C terminal tail anchors of BCL-2 family proteins exert allosteric influence over the interface crucial for BH3 binding and cell survival. This is regulated by additional features taking place at the mitochondria membrane such as recruitment of the death executioner BAX, which, in response to BH3 binding antagonism, contributes to protein complex disruption.

*Summary:* BCL-xL exerts an essential cell survival function which relies on its hydrophobic groove binding to BH3 domain of BH3-only initiators and downstream BAX/BAK executioners. Combining resonance energy transfer assays and molecular dynamics simulations, we unravel that the C-terminal tail mediated subcellular membrane anchoring of BCL-xL selectively advantages binding to membrane-anchored PUMA initiator over BH3 mimetic ligands of the groove. This is due to the combined allosteric effect on BH3-in-groove binding of BCL-xL and PUMA tail anchors. Moreover, doubly anchored PUMA / BCL-xL complexes recruit endogenous BAX, which favors their antagonism by BH3 mimetics. BAX’s C-terminal tail anchor alone is sufficient to enhance BH3 mimetics induced death in cells expressing PUMA / BCL-xL. Thus, the survival function of BCL-xL is regulated by a complex interplay between its tail anchor and those of its interacting partners. This enables both resistance to pharmacological inhibitors and modulation by BAX, which functions as a crucial feedback disruptor of the BCL-xL network.

*Graphical abstract:* 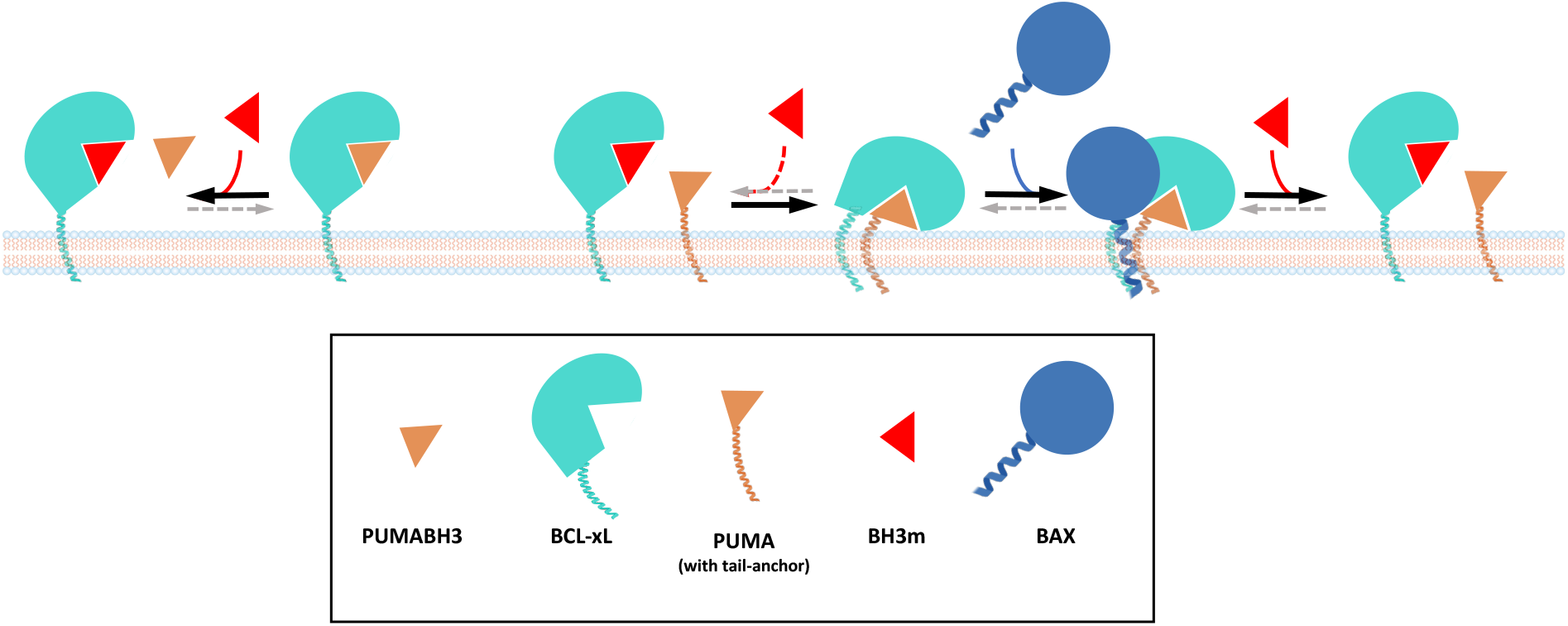

*Highlights:* BH3 mimetic antagonism, and subsequent cell death, are limited when full length BCL-xL binds to some membrane-anchored BH3-only proteins such as PUMA. The BH3-in-groove interface is allosterically modulated by tail anchors of PUMA and BCL-xL. Binding to PUMA enriches BCL-xL interactome and recruits BAX. BAX counteracts the effects of tail anchors in BCL-xL complexes.

## INTRODUCTION

Members of the BCL-2 family engage in a network of interactions to regulate mitochondrial outer membrane permeabilization (MOMP) and subsequent cell death in response to diverse intracellular signals during development and in various diseases. They are classified into three subgroups based on their functions: guardians, which sequester executioners (BAX and BAK), capable of forming pores leading to MOMP, and initiators (BH3-only proteins) which inhibit the function of guardians and additionally activate executioners (Czabotar & Garcia-Saez, 2023). Extensive functional and structural investigations have underscored a conserved binding interface shared among anti-MOMP members (guardians), involved in engagement of the BH3 domain of pro-MOMP counterparts (executioners and initiators) and pivotal in the regulation of cell survival by BCL-2 family proteins.

The structure of the soluble BH3 binding interface of BCL-xL, both unoccupied and in interaction with peptides or chemical compounds, was determined by X-ray crystallography and/or nuclear magnetic resonance (NMR) spectroscopy, revealing that the BH3 domain engages a hydrophobic binding groove of the guardians, formed by α3-α4 with the core helix α5 (Sattler *et al*, 1997; Lee & Fairlie, 2019). These structural insights have accompanied the development of BH3 mimetics (BH3m), small molecules rationally designed to competitively and selectively occupy the groove. BH3m offer substantial therapeutic potential in oncology, as evidenced by the clinical use of venetoclax for the treatment of certain cancers (Juin *et al*, 2013; Diepstraten *et al*, 2022). Optimizing the effects of these compounds is nevertheless essential, as they exhibit dose limiting side effects, and can lead to suboptimal responses to antagonists, which induce partial MOMP, potentially generating of persister cells with enhanced metastatic potential via induction of the integrated stress response (Kalkavan *et al*, 2022) and/or increased genomic instability due to sustained DNA damage (Ichim *et al*, 2015).

Understanding the mechanism of action of BH3m implies to decipher how guardians participate in intracellular interactions within whole cells. Guardians sequester specifically initiators or executioners with different affinities depending on their location and abundance. Intracellular localization of the resulting complexes depends on the identity of the ligands bound (King *et al*, 2022; Beigl *et al*, 2024). Intracellular mobility of BCL-xL is in particular increased by its interaction with BAX and, to a lesser extent, BAK, while it is decreased by interactions with initiators and other proteins not belonging to the family (Todt *et al*, 2015; Vuillier *et al*, 2018; Lauterwasser *et al*, 2021; King *et al*, 2022). This diverse interaction network is also highly dynamic (Kale *et al*, 2018). For example, the release of pro-MOMP factors such as BIM, PUMA or tBID by BH3m molecules antagonizing BCL-xL can be compensated for by their sequestration by MCL-1 to promote cell survival, as observed in solid tumors (Soderquist *et al*, 2018). Consequently, when evaluating the molecular action of BH3m (i.e., disruption of targeted guardian interactions) through cell death induction in cellular or animal models, it is important to note that this method may lack direct information about the antagonistic effects of BH3m molecules. Another level of complexity is provided by the fact that BCL-2 interactions predominantly occur at subcellular membranes (Popgeorgiev *et al*, 2018). All these features indicate that methods preserving cellular integrity, and allowing to compare different interactions between full length proteins are indispensable to evaluate BCL-2 network assembly dynamics. BRET or FRET (Bioluminescence/Fluorescence Resonance Energy Transfert) approaches to monitor the interactions of BCL-2 family members in whole live cells have been used to evaluate the impact of post-translational modifications of BCL-xL response to BH3m (Bah *et al*, 2014) and to observe notable differences in BCL-xL response to BH3m, depending on the pro-apoptotic ligand it engages, from that observed in cell-free studies with recombinant proteins (Aranovich *et al*, 2012; Osterlund *et al*, 2022; Pécot *et al*, 2016).

Interactions between members of BCL-2 family at subcellular membranes enhance their affinity and modulate the hierarchy of interactions between family members (Lovell *et al*, 2008; García-Sáez *et al*, 2009; Yao *et al*, 2015; Chi *et al*, 2019; Yang *et al*, 2019). Most members of the BCL-2 family harbor a hydrophobic *α* helix in their C terminus (hereinafter named tail anchor, noted TA), which allows insertion primarily into the outer mitochondrial membrane or endoplasmic reticulum membrane (Nechushtan *et al*, 1999; Kaufmann *et al*, 2003; Wilfling *et al*, 2012 ; Andreu-Fernández *et al*, 2016). These domains in executioners and initiators are required for their optimal pro-MOMP activity (Zhang *et al*, 2016; Stehle *et al*, 2018; Chi *et al*, 2019). The corresponding domains in guardians contribute to their anti-MOMP activity (Nguyen *et al*, 2024) and are also involved in functions beyond membrane addressing: the MCL-1 TA is implicated in its post-translational regulation, BCL-xL TA contributes to the retrotranslocation of executioners and to the allosteric regulation of initiator proteins in BCL-xL complexes for BAX activation (Todt *et al*, 2013; Bogner *et al*, 2020). Recent studies indicate that direct interactions between the TAs of BCL-2 family proteins also contribute to the regulation of apoptosis (Zhang *et al*, 2016; Andreu-Fernández *et al*, 2017; Lucendo *et al*, 2020; García-Murria *et al*, 2020; Beigl *et al*, 2024; Nguyen *et al*, 2024). It is noteworthy that the BCL-xL TA determines the hierarchy of interactions within the membrane-anchored BCL-2 family members (Bleicken *et al*, 2017). We showed that membrane anchoring of BCL-2 and BCL-xL limits the antagonistic action of BH3m, but the underlying mechanism had remained mostly uncharacterized so far (Pécot *et al*, 2016).

One limitation hindering a comprehensive understanding of how these proteins regulate MOMP is that structures of BCL-2 family protein complexes in the subcellular membrane, their primary functional site, are not well-determined. Evidence from cellular biochemical analysis of higher-order molecular complexes with BCL-xL or MCL-1 challenges the notion that guardian control pro-MOMP partners solely by forming heterodimers (Singh *et al*, 2017; Yang *et al*, 2019; Bogner *et al*, 2020). Moreover, the dynamic nature of complex assembly needs to be taken into account. The conformational dynamics of BCL-2 family proteins manifest through allosteric regulation, wherein distant modifications induce conformational changes that modulate the protein’s activity. For instance, allosteric regulation of BCL-xL controls BAX activation, p53 or initiator sequestration (Follis *et al*, 2018; Bogner *et al*, 2020). In the case of binding to initiators, PUMA or BIM are in fact intrinsically disordered proteins, where the intrinsically disordered region (IDR) of the BH3 domain adopts an amphipathic α-helical conformation upon binding to guardians (Hinds *et al*, 2007; Rautureau *et al*, 2010) and BH3 domain helical stability contributes to affinity (Modi *et al*, 2013). Structural and molecular dynamics (MD) studies also highlighted a high degree of conformational diversity in BCL-xL itself, with the engagement of BH3 peptides inducing significant conformational changes in *α* helices, particularly at the BH3-in-groove interfaces (Yang & Wang, 2011; Liu *et al*, 2015; Liu *et al*, 2017; Lee & Fairlie, 2019). The hydrophobic binding groove of BCL-xL represents a so-called cryptic binding site of remarkable flexibility which can be transiently exposed when unbound (Mizukoshi *et al*, 2020), but that turns into a stably open binding site with different conformations upon engagement by BH3 domains or mimetics, according to an induced-fit mechanism (Rajan *et al*, 2015; Bekker *et al*, 2021; Bekker *et al*, 2023). Binding of the PUMABH3 peptide results in the most significant conformational modification of BCL-xL, α2 and α3 becoming highly disordered with potential functional consequences (Follis *et al*, 2013). While it seems plausible that membrane anchoring might somehow constrain such dynamic conformational changes during protein assembly, whether, how and with which biological consequences it might do so has remained mostly unexplored. As membrane proteins are difficult to study experimentally at the molecular level (White, 2023), MD simulations permit to simulate what could appear in biological context, especially for binding interfaces characterized by significant conformational dynamics. In particular, these studies have unveiled structural disparities between soluble and membrane-anchored BCL-xL, as well as the dynamics of BH3-in-groove positioning concerning the membrane (Maity *et al*, 2013; Ryzhov *et al*, 2020). MD simulations also enabled energetic analysis, which highlighted new determinants of specificity and affinity in the interaction network of the BCL-2 family (Ivanov *et al*, 2016).

Based on these premises and incognita, we herein investigated how BCL-xL TA modulates its protein binding activity in an integrated cellular context: we evaluated dose-response analysis of BRET signals between a wide range of initiator and BCL-xL variants and performed biochemical and proteomic analysis of BCL-xL complexes in cells. Additionally, we performed molecular dynamics simulations and MM/GBSA analysis (Molecular Mechanics/Generalized Born Surface Area) to model different BCL-xL complexes in membrane in order to infer the influence of TA of each binding partner on complex formation. Through this unprecedentedly comprehensive description of the interplay between BCL-2 proteins’ TAs and BH3 binding, we show that the former allosterically restrict antagonism of the latter by BH3m and subjects membrane-anchored BCL-xL binding to negative feedback by the BAX executioner.

## RESULTS

### In whole cell configuration, BCL-xL binds to BH3 domains via the canonical mode

We investigated whether the robustness of certain BCL-xL interactions with BIM and PUMA in response to BH3m antagonism in a whole-cell context is due to a binding mode that is mechanistically distinct from the canonical one (that is, involving the groove occupied by a BH3 helix). We investigated by BRET assays interactions between full length BCL-xL used as an acceptor (_Y_BCL-xL fused to Yellow Fluorescent Protein, indicated by a subscripted Y) with a range of 36 mer peptides comprising the BH3 domains of PUMA, BIM and tBID used as donors (_R_PUMABH3, _R_BIMBH3 and _R_BIDBH3 fused to Renilla Luciferase indicated by a subscripted R) (Supplementary Table 1). BRET donor saturation assays indicated specific interactions in all cases (Figure 1A & Extended description). Within these BH3 domains, we substituted the amino acids engaged in the interaction within the BH3-in-groove of guardian proteins at the H2 and H4 positions, along with the conserved aspartate residue, with alanine (BIM 2A, tBID 3A, PUMA 3A, Supplementary Table 1). As expected, we found that the resulting variants lost specific binding to BCL-xL in BRET assays (Figure 1A). The BCL-xL-specific BH3m compounds (WEHI-539, A-11555463, A-1331852) exhibited dose-dependent antagonism of BH3 domain / BCL-xL interactions (Figure 1B). Their efficacy coincided with their published affinities for BCL-xL BH3 binding pocket, with WEHI-539 being less efficient than A-1155463, itself less efficient than A-1331852 (Tao *et al*, 2014; Leverson *et al*, 2015; Osterlund *et al*, 2022). These results argue that, when BH3 domains are excised from the context of the whole proteins (and from that of PUMA or BIM in particular), they bind to BCL-xL groove in a canonical manner, as described by multiple structural studies using an acellular model. This implies that structure activity relationships of the BH3 binding pocket of BCL-xL are comparable in cell free assays or in intact cells, and that BH3m occupy this pocket in similar manners regardless of contexts.

**Figure 1:**
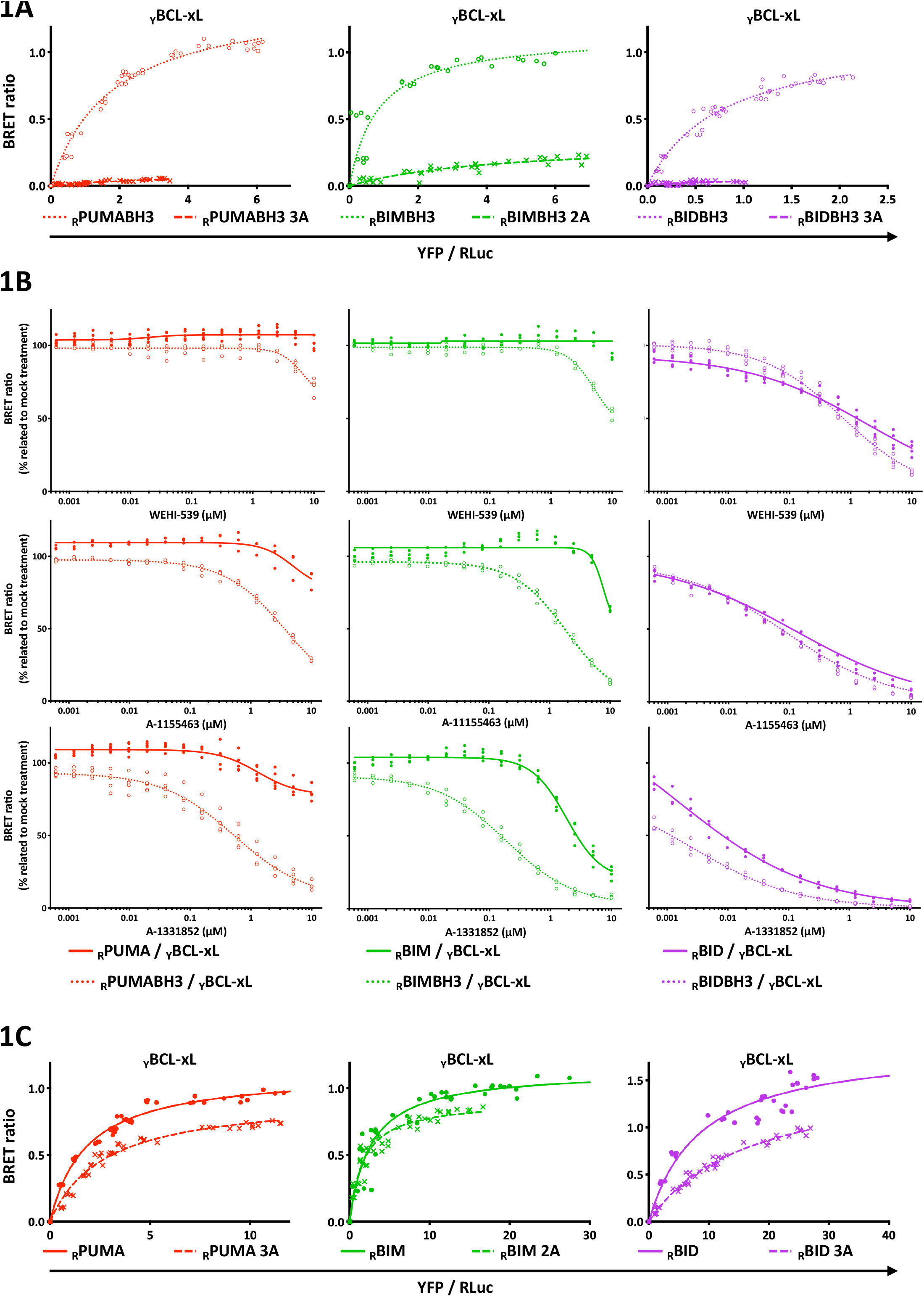
Comparative analysis of the binding of initiators and their BH3 domains to BCL-xL and antagonism with BH3m molecules. 1A-Involvement of hydrophobic amino acids H2, H4 and conserved ASP of PUMA, BIM and tBID BH3 domains in binding of BCL-xL. BRET donor saturation assays were conducted using _R_BH3 domain (empty disk mark), both fused to RLuc, and _Y_BCL-xL fused to YFP in MCF-7 cells. The variant with ALA substitutions (cross mark) (Supplementary Table 1) was compared to the wild-type variant. Data were fitted with a non-linear regression equation assuming a single binding site using GraphPad Prism version 8 for macOS. The data presented are representative of at least two independent experiments. 1B-Dose-response analysis of three BH3m (WEHI-539, A-1155463 and A-1331852) selectively antagonizing _Y_BCL-xL, interacting with either initiator or its BH3 domain alone fused to RLuc in MCF-7 cells. Each BRET signal is normalized to the untreated condition (100%). The graphs include data points from at least three independent replicates. Some of the marks may be superimposed on the graph. All data were fitted using an equation: [Inhibitor] vs. response - Variable slope using GraphPad Prism version 8 for macOS. 1C-Involvement of hydrophobic amino acids H2, H4 and conserved ASP of PUMA, BIM and tBID in binding of BCL-xL. BRET donor saturation assays were conducted using _R_BH3-only initiator (full disk mark) and _Y_BCL-xL in MCF-7 cells. The variant with ALA substitutions (cross mark) (Supplementary Table 1) was compared to the wild-type variant. The representation follows the description provided in Figure 1A.

### BH3-in-groove interactions are non-unique yet necessary determinants for the binding of full length BCL-xL to initiator proteins in whole cells

BRET signals between initiator proteins and the BCL-xL G138E R139L I140N variant, carrying three mutations in residues critical for BH3 binding, were shown to be reduced (Pécot *et al*, 2016). A robust set of additional evidence further argued that protein complexes formed in whole cells between BCL-xL and full length executioners rely, albeit not solely, on BCL-xL canonical BH3 binding site. Firstly, the ALA substitutions in the BH3 domain (PUMA 3A, BIM 2A and tBID 3A, Supplementary Table 1) impacts the saturation curve, although this effect is less pronounced for PUMA 3A and BIM 2A compared to their BH3 domain alone (Figure 1C). Secondly, BRET assays showed that PUMA or BIM interactions with BCL-xL were antagonized by the more affine BH3m A-1155463 or A-1331852 (Figure 1B), even though the concentrations required were much higher than these required to displace interactions with BH3 domains alone. These compounds also inhibited BAD and tBID interactions with BCL-xL in BRET assays and were more efficient to do so than WEHI-539 (Figures 1B & S1A). To control the specificity of these assays, we checked that neither BCL-2 specific ABT-199 nor MCL-1 specific S-63845 affected BAD / BCL-xL interactions, despite inhibiting BAD / BCL-2 and BAK / MCL-1 interactions respectively (Figures S1A & S1B). *A contrario*, the former interactions were neither inhibited by specific BH3m of other guardians. Thus, the effects of BH3m in whole cells reflect their relative affinities and selectivities for BH3-in-groove interfaces of BCL-xL determined in cell free assays (Chen *et al*, 2005; Ivanov *et al*, 2016).

Thirdly, we substituted conserved GLY or ALA residues (A145V, G96V and G35V of PUMA, BIM and tBID, respectively, Supplementary Table 1) to perturb the flexibility of the helix in their BH3 domains, a parameter that affects their affinity in the hydrophobic groove (Day *et al*, 2008; Modi *et al*, 2013; Sora & Papaleo, 2022). Although this substitution modestly affected BRET donor saturation assays, it notably increased sensitivity to WEHI-539 (Figures S1C & S1D).

Overall, these experiments are consistent with evidence from the BH3 peptides, indicating that the formation of BCL-xL complexes with initiators in whole cells is guided by, although not exclusively, BH3 binding, in a manner dependent on the *α* helix structure and its composition in amino acids. However, the robustness of PUMA or BIM interactions with BCL-xL to BH3m antagonism, compared to the BH3 domain alone, suggests that additional molecular features endow these initiator proteins with a competitive advantage over BH3m for binding to BCL-xL.

### The C terminal end of PUMA and BCL-xL cooperate to form BH3 mimetic resistant complexes

Regardless of the BH3m used, PUMA or BIM complexes with BCL-xL consistently exhibited higher resistance to antagonism compared to those with tBID or BAD (Figures 1B & S1A). These differences in sensitivity follow the same hierarchy as this reported by D. Andrews’s laboratory in whole cell FLIM-FRET assays (Liu *et al*, 2019; Osterlund *et al*, 2022). A notable difference was that we observed a greater sensitivity of tBID complexes compared to BAD ones. We ascribe this to the fact that these studies employed mouse tBID whereas we used the human protein. In BRET assays, the former formed complexes with BCL-xL that were less sensitive compared to the latter (Figure S1E).

We focused on PUMA / BCL-xL interactions and searched for molecular features that might provide a competitive advantage to PUMA over BH3m. We designed a chimeric protein in which the PUMABH3 domain was replaced with that of BID (PUMA-BIDBH3, Figure 2A & Supplementary Table 1), whose interaction with BCL-xL is the most sensitive to BH3m antagonism (Figure 1B). The interaction of this domain-swapped chimera with BCL-xL, unlike that of tBID, was resistant to WEHI-539 and significantly less sensitive to A-1131852 (Figures 2B & S2A). Thus, domains surrounding the PUMABH3 domain contribute to enforce interaction with BCL-xL. Of note, the G145V substitution (corresponding to the G35 residue in tBID) in PUMA-BIDBH3 resulted in sensitivity to WEHI-539, similar to what was observed for tBID (Figure S2B), indicating that a BH3 binding mode was still contributing to engagement of the chimeric protein. Deletion of the C-terminal end from PUMA (annotated PUMAdC and PUMA-BIDBH3dC) sensitized their interaction with BCL-xL to antagonism by BH3m (Table Supplementary Table 2, Figures 2B, 2C & S2A). Deletion of the N-terminal end of PUMA (annotated dNPUMA and dNPUMA-BIDBH3) had no detectable effect on BH3m sensitivity (Figures 2B & 2C). These results show that the C-terminal domain of PUMA confers a selective advantage over BH3m for binding to BCL-xL. Of note, deletion of the TA of BCL-xL (BCL-xLdC, Supplementary Table 2) did not enhance BH3m sensitivity of PUMAdC / BCL-xL BRET signals as markedly as it did for PUMA / BCL-xL ones (Figure S2C). Likewise, we observed no striking difference to BH3m treatment between BCL-xL or BCL-xLdC interactions with the sole BH3 domains of PUMA (or BIM or BID, Figure 2D). These results put forth a functional cooperation between the C terminal end of PUMA and that of BCL-xL to promote robustness against BH3m induced displacement.

**Figure 2:**
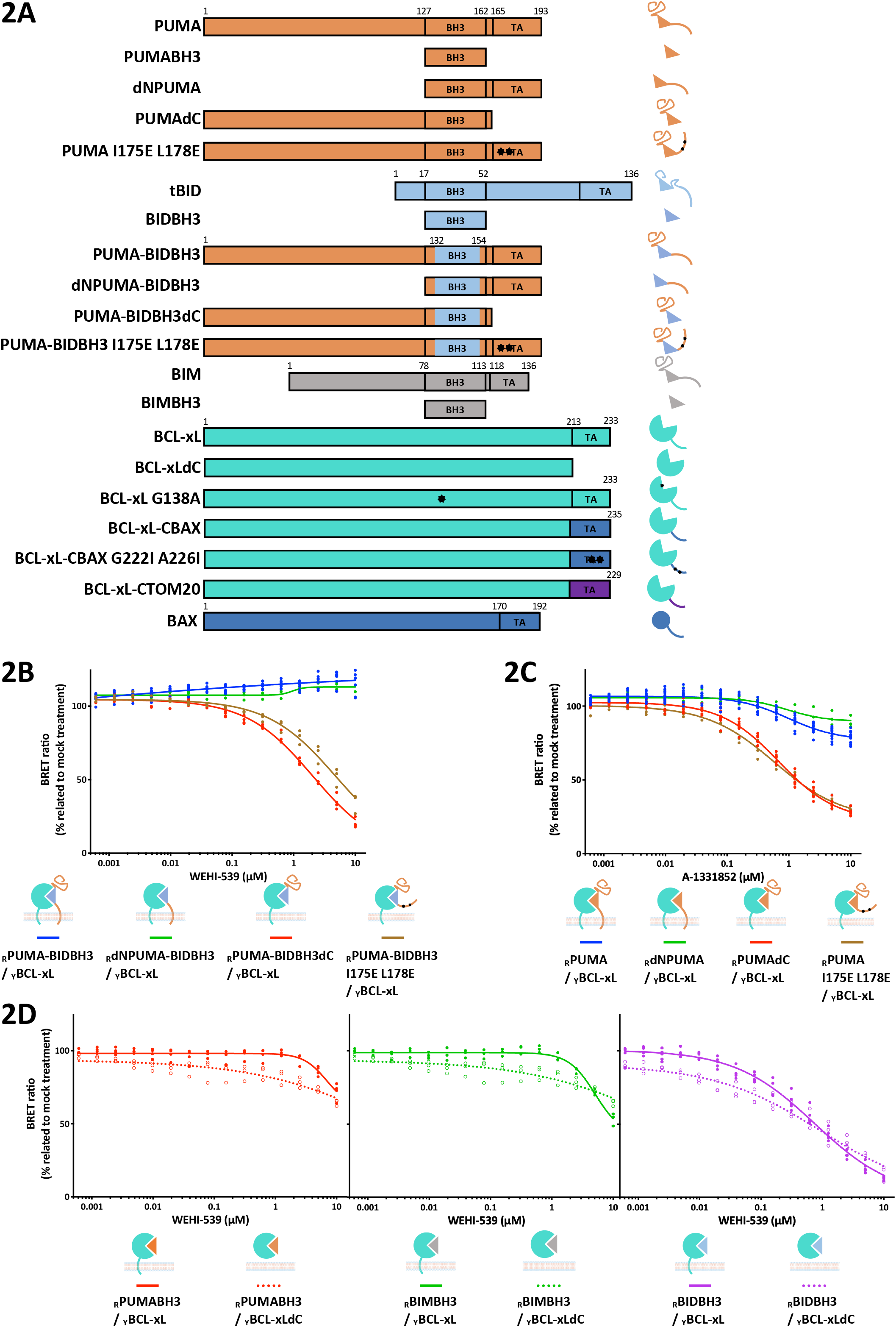
PUMA and BCL-xL tail anchors cooperate to limit BH3 antagonism. 2A-Schematic representation of different protein variants expressed in the cells for this study. BH3 domain of initiators and TA are indicated. The numbering indicates the amino acid position in the full length protein. The pictograms used to describe the complexes evaluated in the BRET dose responses analysis are displayed on the right. See Supplementary Table 1 for sequences. 2B-Dose-response antagonism of _Y_BCL-xL by WEHI-539, depending on the interaction with _R_PUMA-BH3BID variants in MCF-7 cells. The representation is as described in Figure 1B. 2C-Dose-response antagonism of _Y_BCL-xL by A-1331852, depending on the interaction with _R_PUMA variants in MCF-7 cells. For ease of comparison, the data for the response of the _R_PUMA / _Y_BCL-xL complex shown in Figure 1B have been reproduced. The representation is as described in Figure 1B. 2D-Dose-response antagonism of _Y_BCL-xL and _Y_BCL-xLdC by WEHI-539 interacting with _R_BH3 domains of initiators in MCF-7 cells. For ease of comparison, the data for the responses of the _R_BH3 domain / _Y_BCL-xL complexes shown in Figure 1B have been reproduced. The representation is as described in Figure 1B.

### Membrane anchoring of PUMA and BCL-xL constrains orientation of the BH3-in-groove interface towards the lipid membrane

The C-terminal TA sequences of PUMA and of BCL-xL mediate their targeting to mitochondrial and endoplasmic reticulum membranes, in an alkaline-resistant manner demonstrating stable membrane anchoring (Kaufmann *et al*, 2003; Wilfling *et al*, 2012). We confirmed that a variant of PUMA lacking this domain, named PUMAdC and fused to mCherry, exhibited a ubiquitous localization upon expression MCF-7 cells (Figure S2D). As expected, I175E and L178E substitutions in the PUMA TA loose subcellular membrane targeting (PUMA I175E L178E Supplementary Table 2, Figure S2D)(Pemberton *et al*, 2023). Loosening of BCL-xL anchoring to subcellular membranes by the A221R substitution in this TA resulted in variant interacting with PUMA with increased sensitivity to BH3m antagonism (Pécot *et al*, 2016). Similarly, these substitutions, in either native PUMA or chimeric PUMA-BIDBH3, also enhanced sensitivity to BH3m of BRET signals with BCL-xL (Figures 2B & 2C). This argues that the membrane anchoring property of PUMA TA contributes to resistance of PUMA-BCL-xL complexes to BH3m antagonism.

To investigate how membrane anchoring of BCL-xL and PUMA might influence BH3-dependent complex formation, we conducted molecular dynamics simulations. Two model conformations, BCL-xL interacting with a PUMABH3 domain, and BCL-xL interacting with dNPUMA, all anchored in a palmitoyl oleyl phosphatidylcholine membrane (POPC), were constructed and compared. Since we observed that the N-terminal part upstream of the BH3 domain of PUMA did not influence the BH3m antagonism (Figures 2B & 2C), we decided to model PUMA starting from its BH3 domain to the C-terminal sequence (130-193). In both models, the BH3 domain was observed to be nestled between the hydrophobic groove of BCL-xL and the upper layer of the membrane, with a similar initial orientation relative to the membrane (Figure 3A). The PUMA linker, bridging its TA to the BH3 domain, was positioned beneath the globular domain of BCL-xL. The MD simulations of PUMABH3 or dNPUMA with BCL-xL complexes over a period of 2000 ns reveals distinct conformational dynamics (Supplementary Movies 1 & 2). In both simulations, the BH3 domain of PUMA always remained tightly bound to BCL-xL, however, the globular part of BCL-xL adopted different orientations relative to the membrane (Figure 3A). Greater freedom of movement was observed in the PUMABH3 / BCL-xL complex, causing the BH3 domain to lose contact with the membrane and become more exposed to the solvent (Figure S3A). To further characterize these differential dynamics, we computed two angles (roll and tilt) to describe the position of the BCL-xL groove relative to the membrane (Figure 3B). Net differences were observed depending on the presence or absence of the PUMA TA. In the dNPUMA / BCL-xL simulation, the roll and tilt angles remained globally stable throughout the dynamic range. In contrast, the PUMABH3 / BCL-xL simulation showed a drastic change around 500-600 ns, corresponding to the exposure of the BH3 domain to the solvent. After 600 ns, significant differences in tilt and roll were observed: 76° ± 8 and 114° ± 11 for PUMABH3 / BCL-xL (over the 600 – 2000 ns interval), respectively, compared to 132° ± 7 and 96° ± 5 for dNPUMA / BCL-xL. These results indicate that anchoring PUMABH3 via its TA affects the orientation of the BH3-in-groove relative to the membrane.

**Figure 3:**
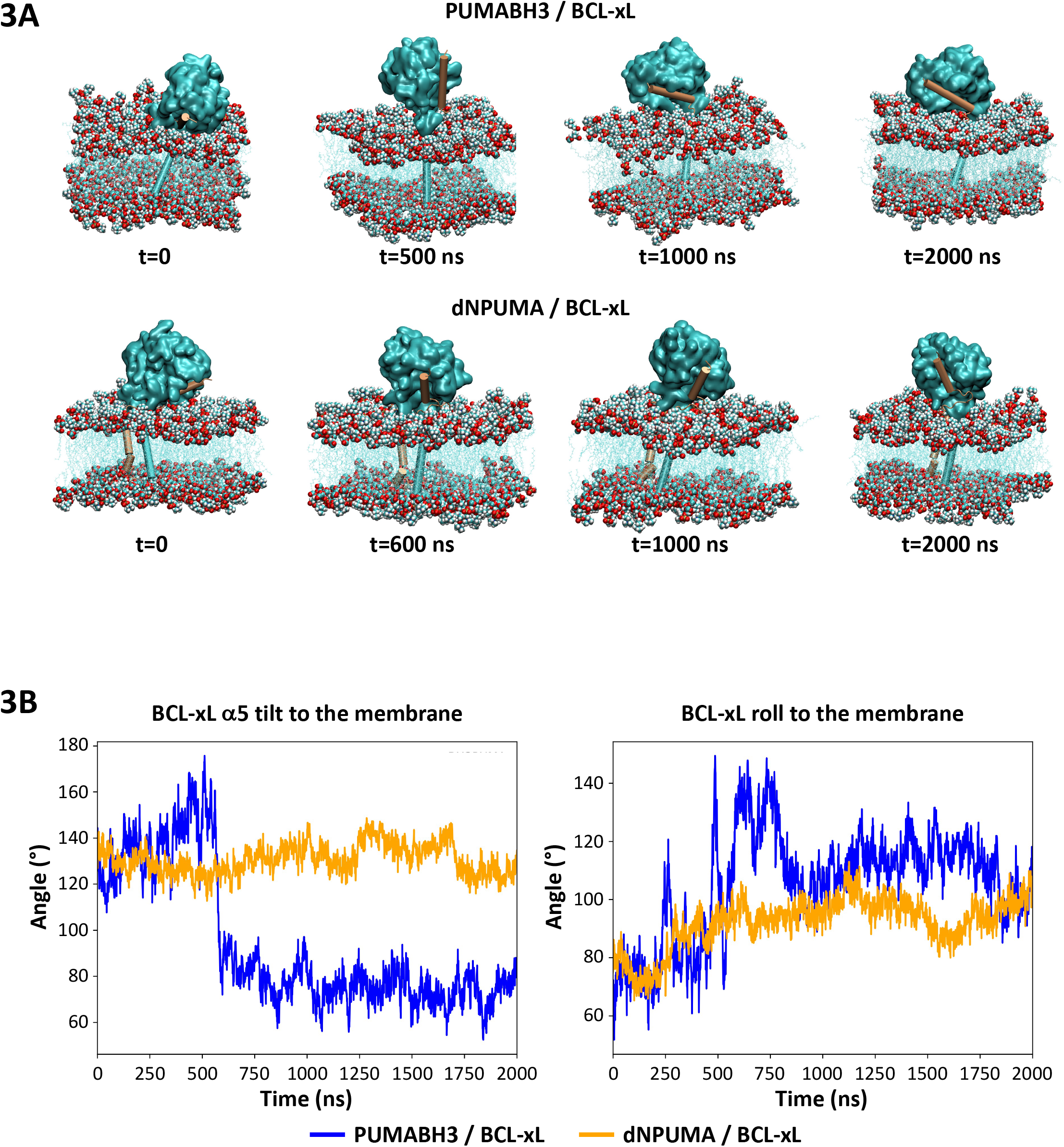
Molecular dynamic simulations analysis of PUMABH3 / BCL-xL and dNPUMA/ BCL-xL complexes. 3A-Representative snapshots from the supplementary movies 1 and 2 illustrating the results of molecular dynamics simulations over time performed to study the behavior of BCL-xL complexes interacting with PUMABH3 or dNPUMA. Phosphatidyl choline headgroups are represented in van der Walls representation and lipid tails are displayed in blue lines to provide a clear view of TA. The helical structure of TA is depicted in Cartoon representation. The simulations were carried out for 2 microseconds. At t=0, the initial orientations of the PUMABH3 / BCL-xL and dNPUMA / BCL-xL complexes are depicted. During the simulation, the PUMABH3 / BCL-xL globular part moves away from the membrane, while this phenomenon was less pronounced for the dNPUMA / BCL-xL complex. 3B-Dynamics of the globular part of the PUMABH3 / BCL-xL and dNPUMA / BCL-xL complexes relative to the membrane over a 2000 ns timespan. BCL-xL tilt was defined as the angle between the BCL-xL α5 helix and the membrane plane (left panel). BCL-xL roll was defined as the angle between a plane formed by BCL-xL α5 and α6 helices and the membrane plane (right panel).

We then focused on the dynamic conformation of BCL-xL TA alone (in PUMABH3 / BCL-xL complex) or in close proximity to the PUMA TA (in dNPUMA / BCL-xL complex). To avoid bias in our modeling procedure, we first determined the initial relative orientation of isolated BCL-xL TA and dNPUMA TA using protein-protein docking methods. In the dNPUMA / BCL-xL complex, both TAs were predicted to have a nearly parallel orientation with their N-terminal sides positioned close together nearly parallel to each other (Figure S3B). No specific amino acid pairs could be identified as prominent in directing the interaction. First of all, we characterized the geometry of BCL-xL’s α9 helix during the simulation (Bansal *et al*, 2000). In the absence of PUMA’s TA, the bend of BCL-xL’s TA helix remained stable throughout the simulation. However, in the presence of PUMA’s TA, the bend increased, suggesting that it constrains the BCL-xL TA helix (Figure S3C). When full length proteins were assembled and virtually heated to room temperature, the PUMA TA helix displayed partial unfolding at the start of the simulation. Then, a kinked conformation first appeared in the PUMA TA helix (t=200ns) and was subsequently adopted by the BCL-xL TA (t=825ns) (Figure S3B). Once this double-kinked conformation formed, both TAs remained close together for the rest of the simulation, with the BH3-in-groove binding interface consistently oriented toward the membrane. In contrast, this kinked helical conformation was never stably observed in the PUMABH3 / BCL-xL complex (Supplementary Movie 1), where the helices remained largely straight, allowing greater freedom of movement for the globular part of the complex and the BH3-in-groove to move further away from the membrane. Remarkably, as the BCL-xL TA becomes more orthogonal to the membrane compared to the start of the simulation, the BH3-in-groove interface moves away from the membrane, as was subtly observed with the dNPUMA / BCL-xL complex (600 ns) and more stably with the PUMABH3 / BCL-xL complex. Overall, our MD studies indicate that the PUMA TA confines the BH3-in-groove interface to the lipid membrane by restricting the dynamics of the BCL-xL TA.

### Double tail anchoring allosterically controls the BH3-in-groove interface

To determine whether and how the TAs of BCL-xL and PUMA might regulate BH3 binding, we performed interaction energy profiling. We calculated the free energy of ligand binding to proteins using the Molecular Mechanics with Generalized Born and Surface Area solvation method (MM/GBSA)(Miller *et al*, 2012). Our MD trajectories provided accurate calculations of the enthalpies for dNPUMA or PUMABH3 interacting with BCL-xL, taking into account membrane anchoring, and allowed decomposition on a per-residue basis. We considered an amino acid to be a significant contributor to affinity if its binding energy is below -1.5 kcal/mol. We mapped the mean per-energy contributions for the 600-2000 ns interval of stimulation onto surface complexes and primary structures of BCL-xL and its ligand PUMABH3 or dNPUMA, during which the complexes exhibited the greatest conformational stability.

As expected, the PUMABH3 / BCL-xL interaction primarily involved residues on both sides of BCL-xL groove (α4 and α5 helices on one side, and the α2 and α3 helices on the other – Figure S4A), which contains the hydrophobic pockets where the hydrophobic residues H1 to H4 of the PUMABH3 domain (I137, L141, M144 and L148 respectively) form van der Waals interactions (Figure S4B). Additionally, the charged amino acids E129, R139 in BCL-xL, along with R135, R142, D146 in the PUMABH3 domain, also contribute to the interaction, consistent with the canonical electrostatic bonds they form (Figure 4)(Ivanov *et al*, 2016; Sora & Papaleo, 2022). Notably, the amino acids F97, E129, L130, N136 and R139 in the BH3 groove of BCL-xL, identified as important for the interaction of the PUMABH3 peptide with recombinant BCL-xL under soluble conditions *in vitro*, are among the identified amino acids with significative energy in our MM/GBSA calculations (Campbell *et al*, 2015). Therefore, anchoring BCL-xL to the membrane per se does not drastically alter the BH3-in-groove interface of the interaction, in agreement with published results (Yao *et al*, 2015).

**Figure 4:**
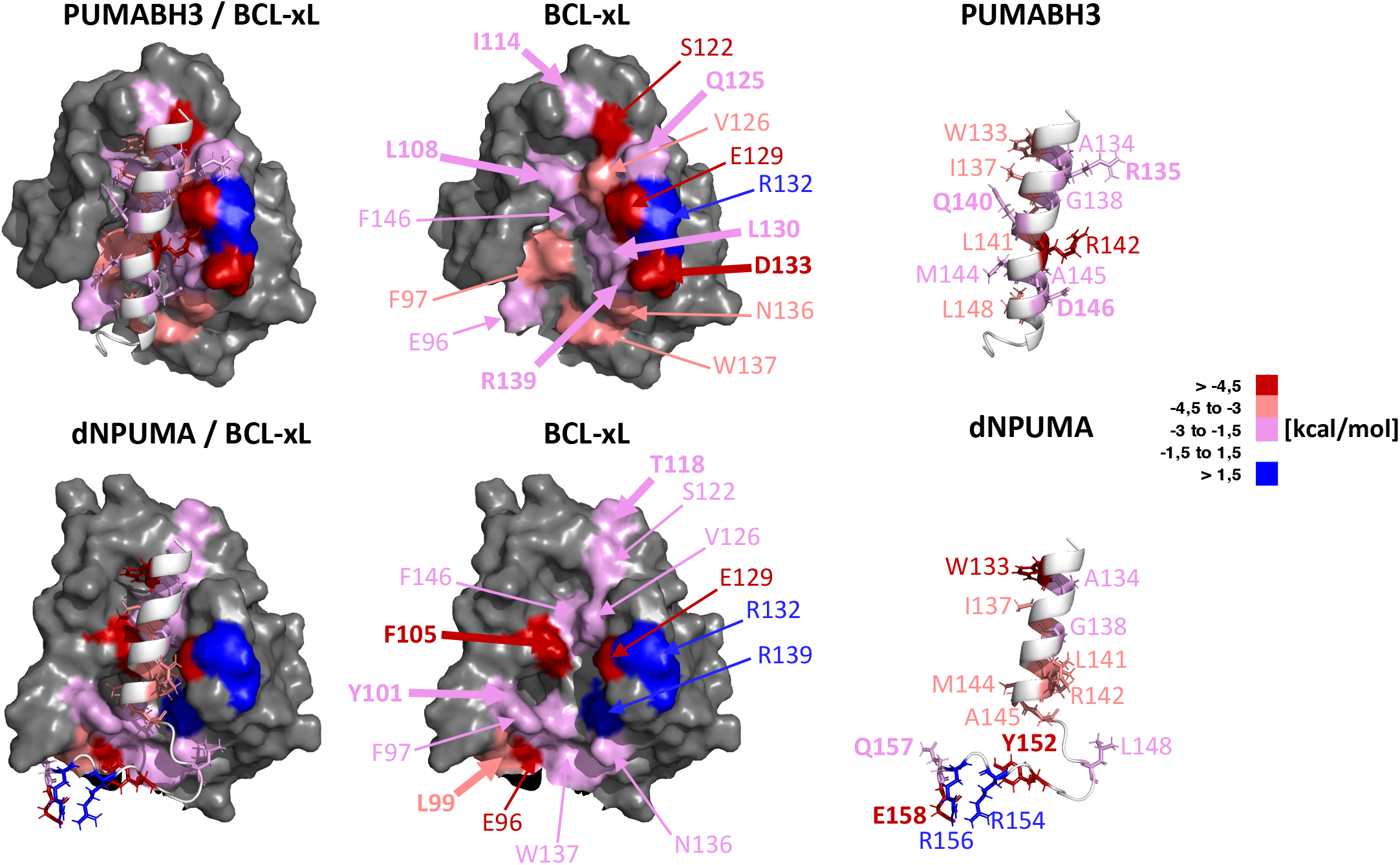
The tail anchors of PUMA and BCL-xL allosterically regulate the BH3-in-groove interface of the PUMABH3 domain within the hydrophobic groove of BCL-xL. PUMABH3 / BCL-xL (upper panels) and dNPUMA / BCL-xL (bottom panels) complexes structures: representative structures (t=1000ns) are colored according to the average per-residue energy contributions, calculated using MM/GBSA from their molecular dynamic trajectories. Only residues with significant binding energy contributions (below -1.5 kcal/mol or above 1.5 kcal/mol) are displayed and annotated according to the color index. Specific residues within the BH3-in-groove of BCL-xL interacting with PUMABH3 or dNPUMA are annotated in bold with a thick arrow (middle and right panels). The underlying numerical values are given in Supplementary Table 3. The C-terminal domains are omitted at the bottom for better readability.

A greater number of BCL-xL amino acids contribute to the interaction with dNPUMA (21 residues) compared to PUMABH3 (14 residues), resulting in a higher overall binding energy for the dNPUMA / BCL-xL complex (-134 kcal/mol) compared to the PUMABH3 / BCL-xL interaction (-80 kcal/mol). Both interactions involve the same key regions (the α2 to α5 helices of BCL-xL and the BH3 domain of PUMA), but the dNPUMA / BCL-xL interaction also includes additional contributions from amino acids in the C-terminal domains of both partners (Figure S4A & S4B). BCL-xL residues (L108, I114, Q125, L130, D133 and R139), which were identified as contributors to the interaction with the PUMABH3 domain (Campbell *et al*, 2015), are not involved in the interaction between dNPUMA and BCL-xL (Figure S4A). Instead, amino acids in the globular part of BCL-xL (L99, Y101, F105 and T118) specifically contribute to the interaction with dNPUMA, increasing the binding surface between BCL-xL and its ligand. On the other hand, the relative involvement of certain residues from BCL-xL (E96, F97, S122, V126, N136, W137) and from the BH3 domain (W133, R142, M144, A145, L148) varies depending on the presence or absence of PUMA TA (Figure 4). To ensure that conclusions were drawn independently from the choice of MM-GBSA parameters, we evaluated the binding energies in relative terms rather than absolute values. The relative contribution of BCL-xL helices varied depending on whether the PUMABH3 domain was anchored in the membrane or not: α2 (12% of the overall binding energy of BCL-xL), α3 (9%), α4 (47%), and α5 (16%) helices with PUMABH3, and α2 (21%), α3 (13%), α4 (20%), α5 (4%) and also α9 (18%) helices with dNPUMA (Figure S4C). As previously reported, the most conserved BCL-xL residues within the BCL-2 family contribute to over 50% of the total binding energy for PUMABH3, but only 26% for dNPUMA (Figure S4C) (Ivanov *et al*, 2016).

Overall, our analysis indicates that PUMA anchoring to the membrane creates a distinct binding site with BCL-xL and introduces additional potential binding interfaces (Figure 4). Notably, the BCL-xL amino acids involved in the binding of BH3m molecules, A-1155463 and A-1331852, exhibit higher binding energy when interacting with dNPUMA compared to PUMABH3 (Figure S4D). This argues that an allosteric regulation of the BH3-in-groove interface by the TA domains of both BCL-xL and PUMA contribute to the differences in competitive BH3m inhibition efficacy observed in our cellular models (Figure 1B).

### Binding to membrane-anchored PUMA stabilizes BCL-xL at subcellular membranes and promotes higher-order complex organization

We investigated the consequences of BCL-xL binding to membrane-anchored PUMA on its intracellular localization and interactome. For this purpose, we used HCT116 and MCF-7 cells expressing bicistronic constructs containing the coding sequence for a flagged PUMA (full length, denoted _F_PUMA or deleted of its TA in C terminal end, _F_PUMAdC) and a green fluorescent protein tagged BCL-xL (denoted _G_BCL-xL), separated by a 2A sequence. This approach allows semi equimolar co-expression of each transgene RNA from a 2A « selfcleaving » peptide sequence (Lopez *et al*, 2016) and biochemical analysis of the complexes from resulting live cells. Engineered cells were obtained by retroviral transduction followed by flow cytometry-based sorting of GFP expressing cells, to ensure comparable expression levels of transgenes across conditions (Figures 5A & S5A). Cells expressing _G_BCL-xL alone, engineered and sorted in a similar manner, were used as controls.

**Figure 5:**
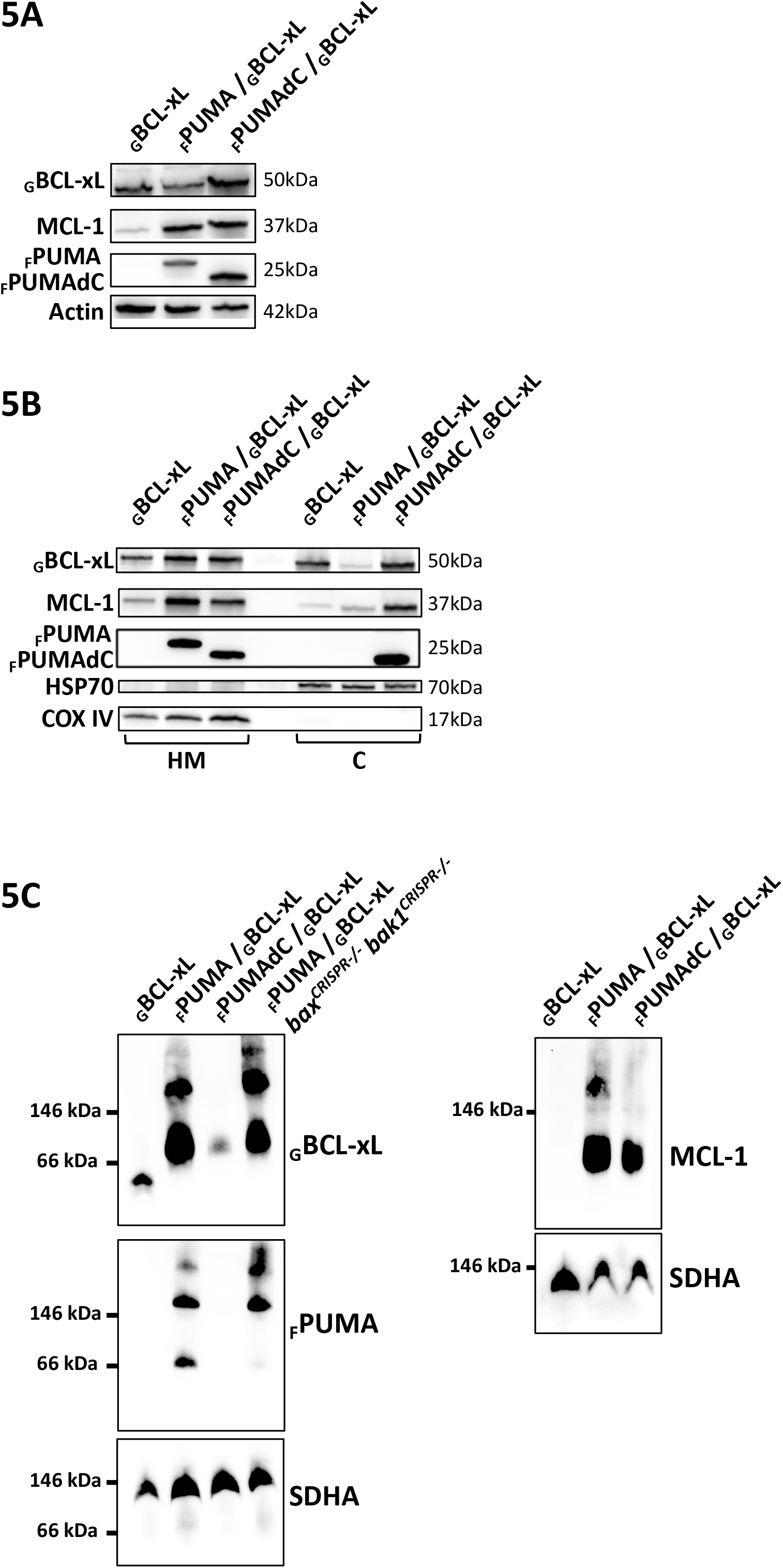
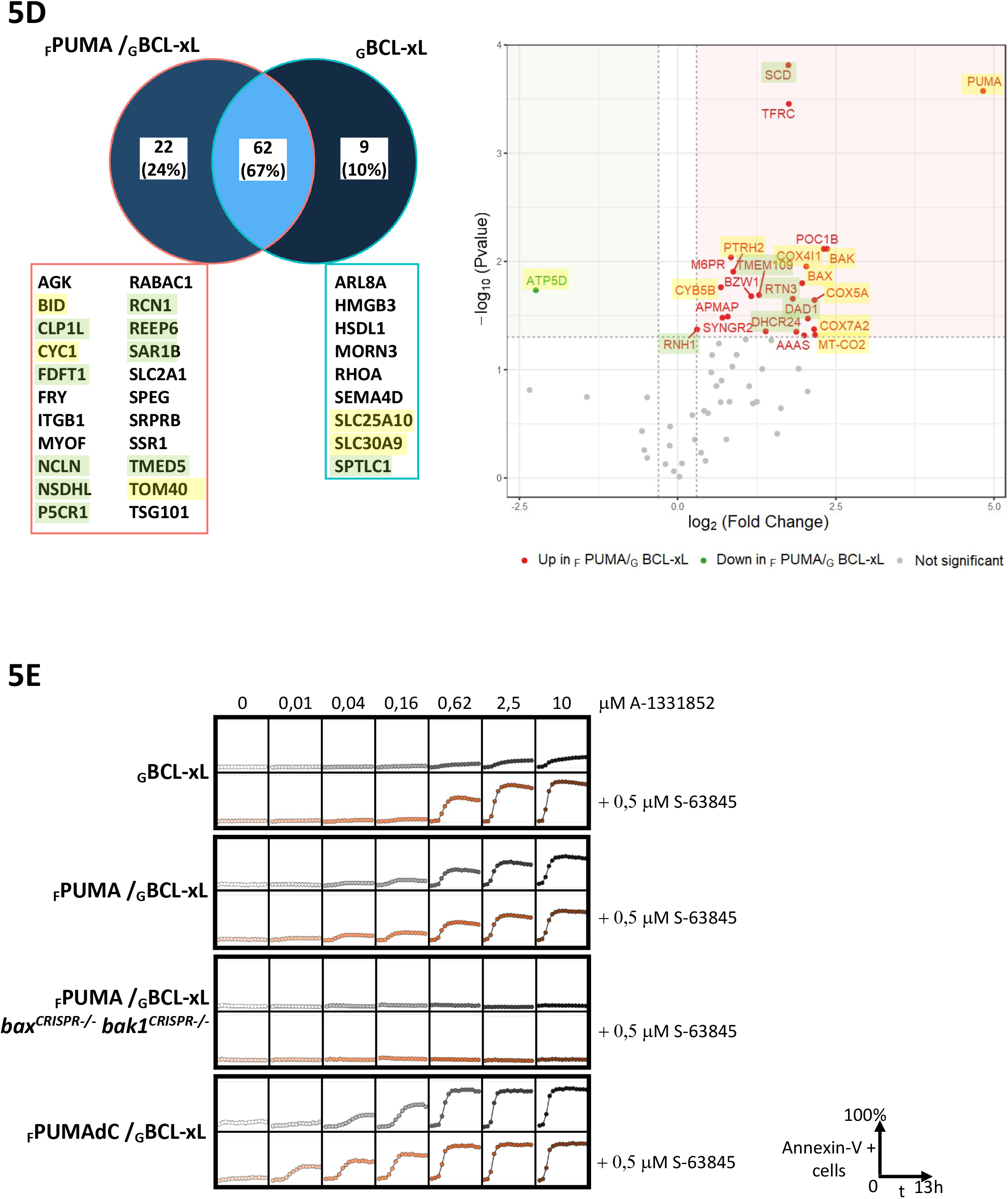
The tail anchor of PUMA stabilizes BCL-xL at the heavy membranes within high molecular weight complexes. 5A-Evaluation of BCL-2 family members expression performed by Western blotting analysis in _G_BCL-xL, _F_PUMA / _G_BCL-xL and _F_PUMAdC / _G_BCL-xL wild-type cells and _F_PUMA / _G_BCL-xL *bax^CRISPR-/-^bak1 ^CRISPR-/-^* HCT116 cells, established in this study. 5B-Subcellular fractionation of BCL-2 family members in heavy membranes (HM, containing mitochondrial and endoplasmic reticulum membranes) and cytosol (C). HSP70 and COXIV serve as controls for cytosolic and mitochondrial markers, respectively. 5C-Analysis of complexes involving _G_BCL-xL, _F_PUMA and/or MCL-1 by blue-native polyacrylamide gel electrophoresis (BN-PAGE) analysis of heavy membranes extracted from _G_BCL-xL, _F_PUMA / _G_BCL-xL and _F_PUMAdC / _G_BCL-xL HCT116 cells. The SDHA protein serves as a loading control. 5D-Comparative analysis of the interactome of _G_BCL-xL in the presence and absence of _F_PUMA in HCT116 cells. Venn diagram illustrating the number of overlapping and specific interactants, that have been detected, in relation to the control, in the interactomes of _G_BCL-xL with and without _F_PUMA. Interactants identified under a single cell line are listed in the red panel for _F_PUMA / _G_BCL-xL and in the blue panel for _G_BCL-xL (left panel). Volcano plot showing the distribution of fold changes in the amount of detected interactants of _G_BCL-xL with and without _F_PUMA. In the plot, red and green markers denote significantly up- and down-interacting proteins, respectively, while gray markers represent non-significantly changed levels of interactants (log2FC>0.3 or log2FC<-0.3 and Pvalue < 0.05) (right panel). Proteins located in the mitochondrial and endoplasmic reticulum membranes are consistently highlighted in yellow and green, respectively, across the entire figure. 5E-Kinetics of single cell apoptosis induction in _G_BCL-xL, _F_PUMA / _G_BCL-xL and _F_PUMAdC / _G_BCL-xL, treated with a concentration range 1:4 serial dilution of A-1331852 (0 to 10 μM) with and without S-63845 (0,5 μM). Involvement of BAX and BAK were evaluated in _F_PUMA / _G_BCL-xL HCT116 cells. Apoptosis was assessed by AnnexinV staining over 13 hours using live cell imaging with the Incucyte S3 imager. The data presented are representative of five independent experiments.

Cellular fractionation of _G_BCL-xL, _F_PUMA / _G_BCL-xL and _F_PUMAdC / _G_BCL-xL HCT116 cells showed that _G_BCL-xL expressed alone distributed between the cytosolic and heavy membrane containing mitochondria fractions (Figure 5B). In contrast, co-expression of _F_PUMA led to an almost exclusive localization of _G_BCL-xL to the heavy membrane fraction, where _F_PUMA was also preferentially localized (also observed in MCF-7 cells, Figure S5B). Coexpression of _F_PUMAdC, which itself co-localized to membrane and soluble fractions, did not enhance the membrane localization of _G_BCL-xL, which was found to be comparable to that observed in cells expressing _G_BCL-xL alone (Figure 5B). Thus, the TA of _F_PUMA enhances the membrane localization of _G_BCL-xL. We then compared protein complexes in membrane fractions from engineered HCT116 cells using native gel electrophoresis (BN-PAGE). In the absence of _F_PUMA, _G_BCL-xL that localized at membranes migrate with an apparent molecular weight corresponding to that of monomers (Figure 5C). Upon co-expression of _F_PUMA, membranous _G_BCL-xL migrated at a molecular weight expected for _F_PUMA / _G_BCL-xL heterodimers but also at higher molecular weights (also observed in membranes fractions of engineered MCF-7 cells, Figure S5C). Membranous _F_PUMA showed a migration profile similar to that of _G_BCL-xL ones. Upon co-expression of _F_PUMAdC, membranous _G_BCL-xL migrated only at the size of heterodimers, indicating that higher-order complex organization requires _F_PUMA membrane anchoring.

To further investigate the nature of these complexes, we performed a pull-down of GBCL-xL from lysates of engineered HCT116 cells using an anti-GFP antibody, prior to mass spectrometry-based identification and quantification of co-immunoprecipitated proteins. We dubbed as potentially aspecific binders and removed from any further analysis: i) proteins that we identified by mass spectrometry in parallel control assays (anti-GFP pull downs from GFP expressing HCT116 cells); ii) proteins identified in a CRAPome (Mellacheruvu *et al*, 2013) with a frequency of more than 15%. After application of this counter-selective filter, 71 and 84 proteins were identified as interactors in _G_BCL-xL and _F_PUMA-2A-_G_BCL-xL HCT116 cells respectively (with 62 in common, 9 and 22 specifically in _G_BCL-xL and _F_PUMA / _G_BCL-xL HCT116 cells, Figure 5D left panel). We identified the mitochondrial outer membranous proteins TOM40 and BID, in addition to vesicle and reticulum endoplasmic proteins, in _G_BCL-xL pull downs only when _F_PUMA was coexpressed. Moreover, fold change analysis revealed that the co-expression of _F_PUMA enhanced interaction between _G_BCL-xL and 23 proteins (Figure 5D right panel), including the multi-domain apoptotic executioners BAX and BAK (investigated further below). A substantial number of these proteins were found to be mitochondrial or endoplasmic reticulum proteins according to gene ontology classifications (Figure S5D).

Altogether, these results argue that engagement to membrane-anchored _F_PUMA enhances the binding of _G_BCL-xL to other mitochondrial partners, assembled into protein complexes larger than _F_PUMA / _G_BCL-xL heterodimers.

### BH3 mimetic antagonism of PUMA / BCL-xL doubly anchored complexes is only partly efficient at inducing cell death, regardless of the compensatory BH3 binding activity of MCL-1

We explored the consequence of double anchoring on cell fate regulation by _F_PUMA / _G_BCL-xL complexes in response to BH3m antagonism. Engineered HCT116 cells described above were challenged with increasing concentrations of the potent antagonist A-1331852 and cell death rates were measured by annexin V staining with time lapse live cell imaging. The coexpression of _F_PUMA constructs with that of _G_BCL-xL significantly sensitized cells to A-1331852 treatment compared to _G_BCL-xL alone, with cell death triggered at lower concentrations of the BH3m, and faster cell death kinetics at high concentrations (Figure 5E). Double invalidation CRISPR of *BAX* and *BAK1* genes in HCT116 coexpressing _F_PUMA and _G_BCL-xL (Figure S5E) abrogated cell death induced by A-1331852 treatment, confirming that BAX and BAK are major executioners of BH3m-induced cell death (Figure 5E). Cells expressing _F_PUMAdC were more sensitive than these expressing _F_PUMA. Coimmunoprecipitation experiments (pulling down the GFP moiety of _G_BCL-xL) confirmed that this differential sensitivity to A-1331852 treatment of _F_PUMAdC versus _F_PUMA binding to _G_BCL-xL coincided with a better displacement of _F_PUMAdC from _G_BCL-xL upon treatment (Figure S5F). Thus, the sensitivity to BH3m antagonism (as measured by BRET analysis and co-immunoprecipitation) and cell death initiation in response to the compound are linked.

We investigated a possible role for MCL-1 (which can engage PUMA) in the relative resistance of _F_PUMA / _G_BCL-xL HCT116 cells. Endogenous MCL-1 expression levels were significantly, and to similar extent, increased in _F_PUMA / _G_BCL-xL and _F_PUMAdC / _G_BCL-xL cells compared to _G_BCL-xL cells (Figure 5A), which we speculatively ascribe to a decrease of MCL-1 proteasomal degradation upon PUMA-BH3 binding (as already described for BIM, Czabotar *et al*, 2007). Endogenous MCL-1 was found to co-immunoprecipitate with _F_PUMA in lysates from engineered cells, regardless of PUMA TA expression (Figure S5F). Subcellular fractionation and BN-PAGE analysis showed that MCL-1 preferentially localized to heavy membranes, as part of complexes of molecular weight higher than that of dimers, when _F_PUMA, but not _F_PUMAdC, was expressed together with _G_BCL-xL (Figures 5B & 5C). Cell death assays described above but this time in the presence of a fixed concentration of S-63845 (0,5 µM) highlighted a synergistic effect of S-63845 combined with A-1331852 in cells expressing _G_BCL-xL alone, confirming that MCL-1 can compensate for BCL-xL antagonism in HCT116 cells by sequestering pro-MOMP factors released by BH3m antagonizing BCL-xL. Strikingly, it also had no detectable effect on the relative resistance of _F_PUMA / _G_BCL-xL cells to BCL-xL antagonism. Addition of S-63845 did not further increase significantly cell death in the sensitive _F_PUMAdC / _G_BCL-xL cells (Figures 5E & S5G). _F_PUMA / _G_BCL-xL cells were induced to die at higher concentrations of A-1331852 combined with S-63845, and the percentage of dead cells was lower than in _F_PUMAdC / _G_BCL-xL cells. Thus, membrane anchoring of PUMA affects the membrane localization and assembly into high order complexes of MCL-1, as it does for BCL-xL, and it limits cell death induced by specific BCL-xL antagonists, even when they are combined with antagonists of MCL-1.

### BAX sensitizes doubly anchored PUMA / BCL-xL to BH3 mimetics antagonism by its C-terminal sequence

Binding partners of BCL-xL in the presence of PUMA identified above may regulate the dynamics of complex assembly and response to BH3m. We focused on BAX and BAK, which we found to be enriched in _G_BCL-xL pull downs upon co-expression of _F_PUMA (Figure 5C). To confirm this enrichment, we performed proximity ligation assays (PLA) between endogenous BAX and _G_BCL-xL in engineered cells (co-expressing _F_PUMA or not, Figure S6A). Flow cytometric analyses were performed to simultaneously quantify BAX / _G_BCL-xL proximity events (marked by PLA fluorescent signals) and _G_BCL-xL expression (marked by GFP fluorescence) in each single cell of _F_PUMA expressing or _F_PUMA free populations. We noted in both cell populations an expected positive relationship between BAX / BCL-xL PLA signals and _G_BCL-xL expression (see Extended description). However, a higher mean intensity and a narrower distribution of PLA signals was found in the _F_PUMA expressing population. Moreover, and most importantly, cells from the _F_PUMA expressing population systematically and significantly exhibited enhanced BAX / BCL-xL PLA signals compared to cells from the _F_PUMA free population expressing the same range of _G_BCL-xL levels (Figure 6A). This observation, made over the whole _G_BCL-xL expression spectrum, indicates that _F_PUMA enhances the proportion of BAX molecules that are proximal to _G_BCL-xL.

**Figure 6:**
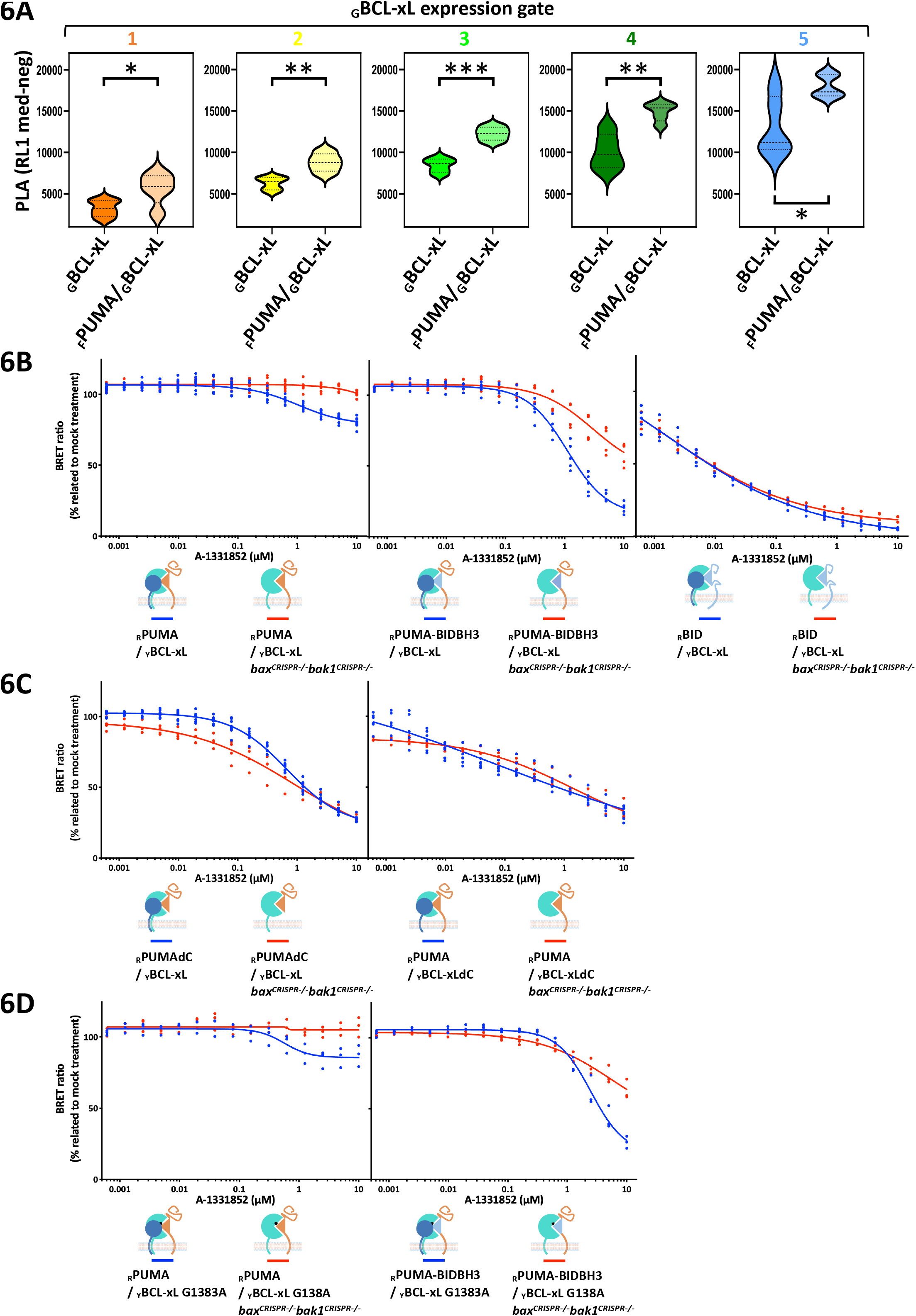
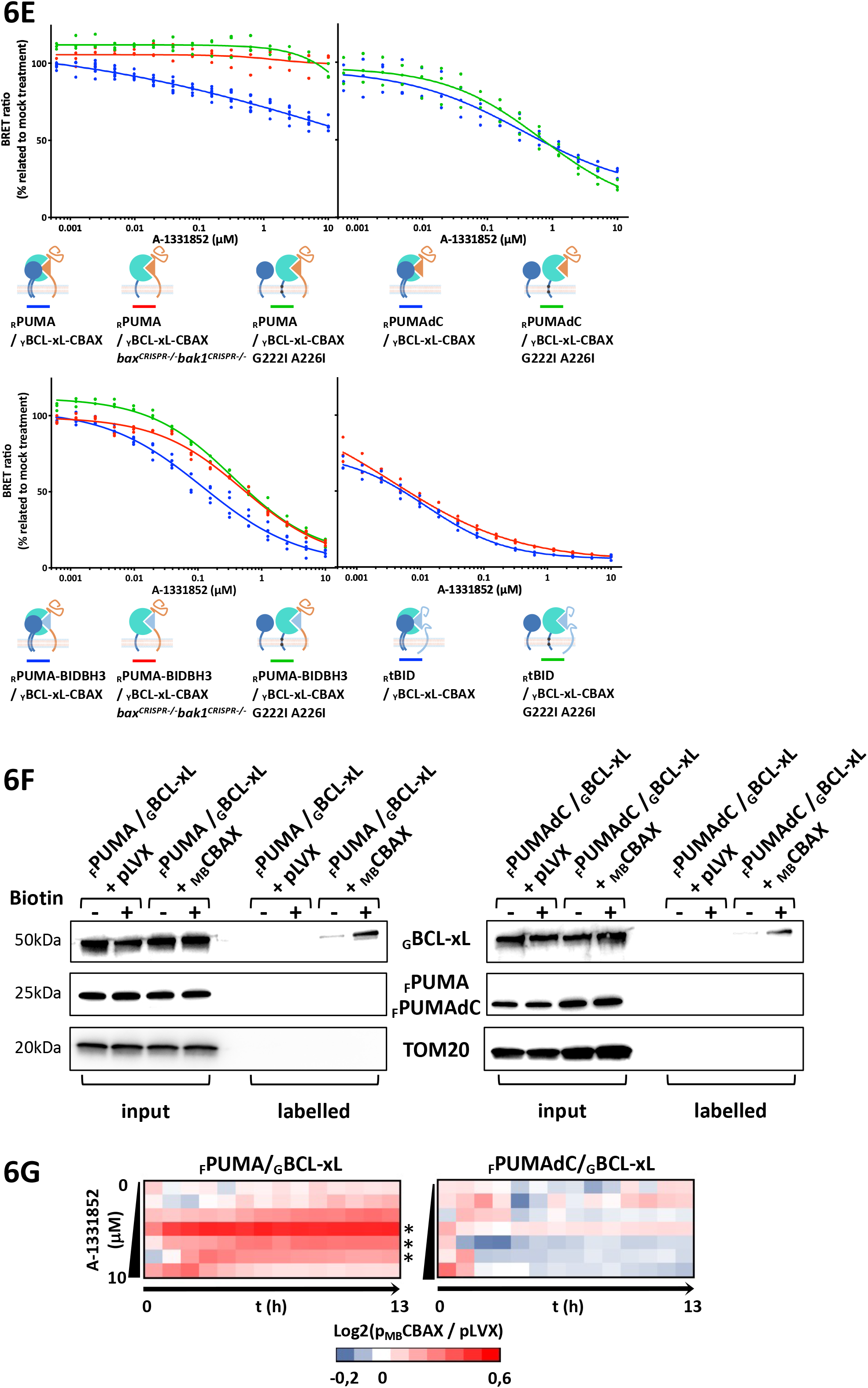
The BAX C-terminal sequence amplifies the antagonistic effect of BH3 mimetics on PUMA / BCL-xL complexes. 6A-Proximity ligation assay of BAX / _G_BCL-xL as a function of PUMA. HCT116 cells expressing either _G_BCL-xL alone or together with _F_PUMA were stained for BAX / GFP interaction. To ensure comparability, five gates with homogeneous _G_BCL-xL expression were defined and PLA signal was analyzed in each rainbow-colored gate (see Figure S8B). The graph illustrates violin plots of the PLA signal (corrected by the PLA in the negative control cells) within gates of _G_BCL-xL expression in three independent experiments. The p-values of the unpaired Student’s t-test between the two cell lines is indicated for each gate. *P < 0.05, **P < 0.01, ***P < 0.001 versus control. Source data are available online for this figure. 6B-6C-6D-6E-Dose-response antagonism of _Y_BCL-xL, _Y_BCL-xLdC_, Y_BCL-xL G138A, _Y_BCL-xL-CBAX, _Y_BCL-xL-CBAX G222I A226I chimeric proteins complexed with either _R_PUMA, _R_PUMA-BIDBH3, _R_tBID, _R_PUMAdC, as a function of BAX and BAK expression. A-1331852 BH3m antagonism of the indicated complexes is compared between wild-type and *bax^CRISPR-/-^ bak1 ^CRISPR-/-^* MCF-7 cells. For ease of comparison, the data for the responses of the _R_PUMA / _Y_BCL-xL, _R_tBID / _Y_BCL-xL, _R_PUMAdC / _Y_BCL-xL, _R_PUMA / _Y_BCL-xLdC, _R_PUMA-BIDBH3 / _Y_BCL-xL complexes shown in Figures 1B, 2C and S2C have been reproduced. The representation follows the description provided in Figure 1B. 6F-Evaluation of the interaction between _MB_CBAX and _G_BCL-xL expressed with _F_PUMA or _F_PUMAdC in HCT116 cells using the BirA *in situ* proximal biotinylation assay. Input and labeled (i.e. biotinylated) proteins isolated with biotin affinity beads were analyzed by Western blotting for _G_BCL-xL, _F_PUMA and TOM20. 6G-Single cell apoptosis analysis in _F_PUMA / _G_BCL-xL and _F_PUMAdC / _G_BCL-xL with or without expression of _MB_CBAX, treated with concentration range 1:4 serial dilution of A-1331852 (0 to 10 μM). Apoptosis was analyzed over 13 hours for AnnexinV staining by live cell imaging using the Incucyte S3 imager. The heatmap summarizes the Log2 ratio of the percentage of cell death in cells expressing _MB_CBAX compared to the control vector (pLVX). Data are mean of five independent experiments. The significant difference in overall kinetics is determined by permutation p-values computed using the comparegrowthcurve function from the statmod R package. *P < 0.05, **P < 0.01, ***P < 0.001

Comparative BN-PAGE analysis of membrane fractions from _F_PUMA / _G_BCL-xL HCT116 cells, either lacking expression of BAX and BAK or not, showed that these executioners are not required for the formation of high molecular weight _F_PUMA / _G_BCL-xL membrane complexes (Figure 5D). In the absence of BAX and BAK, _F_PUMA and _G_BCL-xL proteins consistently formed high molecular weight complexes and tended to produce fewer heterodimeric _F_PUMA / _G_BCL-xL complexes. Thus, when endogenous BAX and BAK are recruited in the proximity of _G_BCL-xL during _F_PUMA expression, they may interfere with _G_BCL-xL, preventing its association with _F_PUMA into multi-molecular complexes.

We then invalidated *BAX* and/or *BAK1* genes by CRISPR in MCF-7 cells to investigate the consequence on BRET signals and their response to BH3m antagonism (Figure S6B). Double invalidation had no drastic effect on the BRET donor saturation assays of BCL-xL interactions with either PUMA, tBID, PUMA-BIDBH3 or BIM (Figure S6C), suggesting that endogenous BAX and BAK do not efficiently compete with the donors for binding to acceptor BCL-xL. However, *BAX* invalidation, and to a lesser extent that of *BAK1*, weakened PUMA / BCL-xL antagonism by BH3m (Figure S6D). The absence of the two executioners completely suppressed the response to A-1331852 treatment of PUMA / BCL-xL complexes (Figure 6B). It also significantly reduced that of PUMA-BIDBH3 / BCL-xL and BIM / BCL-xL, while it had no detectable effect on the sensitivity of tBID / BCL-xL (Figures 6B & S6D). Moreover, *BAX* and *BAK1* invalidation did not promote resistance to BH3m of PUMAdC / BCL-xL or PUMA / BCL-xLdC BRET signals (Figure 6C). Of note, these results imply that the low response to BH3m assessed for PUMA and BIM BRET signals is not due to cell death related loss of donors and/or acceptors signals. Thus, BAX and BAK interferences with PUMA / BCL-xL complexes increase their sensitivity to displacement by BH3m, and it does so only for doubly anchored complexes.

Because BAX and BAK are recruited in the vicinity of BCL-xL upon PUMA co-expression, we sought for BCL-xL binding motifs to these executioners that might be involved in their regulation of PUMA / BCL-xL sensitivity to BH3m. To investigate a role for canonical binding, we used the BCL-xL G138A variant, carrying a substitution in the groove, that renders it defective for binding to BAX and BAK while interaction with PUMA and PUMA-BIDBH3 is retained (Desagher *et al*, 1999; Ding *et al*, 2014; Le Pen *et al*, 2017)(Figure S6E). *BAX* and *BAK1* deletion resulted in decreased BH3m antagonism of PUMA / BCL-xL G138A and PUMA-BIDBH3 / BCL-xL G138A complexes (Figure 6D). Therefore, the effects of BAX and BAK may rely on a BCL-xL interaction site distinct from the groove. This rules out competition with PUMA for binding to this site. The C terminal sequence of BAX interacts with full length BCL-xL and its TA alone in biological membranes (Ding *et al*, 2014; Andreu-Fernández *et al*, 2016). To test whether this non canonical binding contributes to BAX dependent sensitization of double anchored complexes to BH3m antagonism, we first swapped BCL-xL TA with that of the mitochondrial membrane protein TOM20 (Supplementary Table 2), which cannot interact with TAs of BCL-2 family members (García-Murria *et al*, 2020). Double invalidations of *BAX* and *BAK1* genes had no detectable impact on the BH3m response of PUMA / BCL-xL-CTOM20 BRET signals (Figure S6F), in line with the necessity of an interaction involving the BCL-xL TA with that of BAX. In a second set of experiments, we replaced BCL-xL TA with the C-terminal of BAX (Supplementary Table 2). The resulting chimeric protein was shown to robustly associate with the C-terminal sequence of BAX (Andreu-Fernández *et al*, 2016 and see hereafter). The sensitivity to BH3m of BCL-xL-CBAX interaction with PUMA or PUMA-BIDBH3 was enhanced compared to that of BCL-xL (compare Figures 6B & 6E) and lost when the *BAX* and *BAK1* genes were invalidated (Figure 6E). This sensitization was also lost when the G222I A226I substitutions (reported to prevent BAX C-terminal sequences homodimerization, Zhang *et al*, 2016 ; García-Murria *et al*, 2020), were introduced in BCL-xL-CBAX (Supplementary Table 2 & Figure 6E). Note that no effect of this double substitution is observed in the BH3-mimetic dose-response of the complex with PUMAdC (Figure 6E). Thus, modulation of interactions between BCL-xL and BAX TA modify the response to BH3m of PUMA / BCL-xL complexes when and only when they are doubly anchored.

To investigate the impact of the BAX C-terminal sequence on cell death induction by BCL-xL antagonism, we constitutively expressed BAX C-terminal sequence fused to the MYC-labelled biotin ligase BirA (_MB_CBAX) within HCT116 cells co-expressing PUMA / BCL-xL or PUMAdC / BCL-xL (Figure S6G). Biotinylation of _G_BCL-xL was observed (as judged by pull down assays in Figure 6F), indicating that the proximity between the BAX C-terminal sequence and BCL-xL, reported in Andreu-Fernández *et al*, 2016, is maintained when PUMA is co-expressed. No biotinylation of TOM20 and _F_PUMA, both of which are present at the mitochondrial membrane, was detected, suggesting the specificity of the interaction between _G_BCL-xL and BAX C-terminal sequence. Remarkably, expression of the BAX C-terminal sequence resulted in enhanced cell death rates upon BH3m (antagonizing BCL-xL) treatment of PUMA / BCL-xL but not PUMAdC / BCL-xL cells (Figure 6G).

Collectively, these results imply that BAX modulation of the response to BH3m of doubly anchored PUMA / BCL-xL complexes involve interactions between BAX C-terminal sequence and BCL-xL TA.

## DISCUSSION

The present study was initiated based on the observation that in whole cells, membrane localized complexes of BCL-xL with some BH3-only initiator proteins resist to the antagonistic effects of BH3m (Pécot *et al*, 2016). Our results confirm that BH3-in-groove binding, according to molecular motifs characterized *in vitro,* critically determines BCL-xL complex formation in whole cells, and reveal that it is allosterically modulated by TAs of binding partners, endowing initiators with a competitive advantage over soluble BH3m for binding to BCL-xL. This interplay is likely to underlie the reported propensity of membranes and of BCL-xL TA to regulate interaction preferences as shown in a minimal cell free BCL-2 network (Bleicken *et al*, 2017) and it provides a rationale for the heightened pro-survival properties observed for membrane-anchored BCL-xL (Kale *et al*, 2018; Czabotar & Garcia-Saez, 2023). As tBID has been suggested to be membrane-anchored within mitochondrial membranes (Wilfling *et al*, 2012) and engages in BH3m sensitive interactions with BCL-xL, the nature of membrane insertion of the initiator, rather than anchoring *per se*, may play a critical to promote resistance against BH3m antagonism.

BH3 binding relies on an induced-fit mechanism, in which both molecules conformationally adapt to each other (driving for instance binding of BIM BH3 IDR within the cryptic binding site of BCL-xL, as reported by Bekker *et al*, 2023). Thus, novel studies considering conformational dynamics are required to fully apprehend the influence of TAs on this binding in the presence of membranes. We herein performed molecular dynamic simulations of BCL-xL to fill the current gap in this type of study and our analysis brings further support to the notion that the BCL-xL BH3 binding site is highly dynamic, versatile, and capable of adopting alternative conformations (Figure 7). In particular, we unravel an allosteric effect of membrane anchoring on BH3 binding, which manifests at three distinct coordinated levels. Firstly, the binding to membrane-anchored dNPUMA led to maintain the hydrophobic groove of BCL-xL, which is otherwise flexible and transiently exposed to the cytosol as reported for apo-BCL-xL (Ryzhov *et al*, 2020 and our data not shown) towards the membrane. Membrane-oriented confinement of the BH3 binding site may constraint induced-fit BH3 binding as discussed below. Secondly, membrane anchoring of dNPUMA also results in adjustments of the BCL-xL TA within the membrane. Our MM/GBSA analysis suggests potential binding interactions between residues in the C-terminal region of the PUMA BH3 domain and BCL-xL. Although we were unable to provide experimental evidence for direct interactions between the PUMA and BCL-xL TAs alone, we cannot formally rule out the possibility that these may constitute an ancillary binding site to the groove, as proposed in the “double-bolt locks” model (Liu et al, 2019) (Pemberton et al, 2023). Furthermore, while we did not directly associate a specific TA state with BCL-xL’s membrane orientation, our MD analyses showed that the BCL-xL TA helix adopts a kinked conformation in the presence of the PUMA TA, following the transmembrane repositioning of both TAs (Figure 7). Thus, a functional interplay between BCL-xL TA and PUMA TA, which may act as a damper, could influence the membrane-anchored topology of BCL-xL, consequently constraining the dynamics of BH3-in-groove orientation relative to the membrane, otherwise observed in the PUMABH3 / BCL-xL complex. Thirdly, we observed significant differences in the modalities of BH3-in-groove engagement when both BCL-xL and PUMA are membrane-anchored. A thorough comparison of binding energies, with and without the presence of PUMA TA, highlights the involvement of distinct contributing residues within the BCL-xL groove, and reveals changes in the respective contributions of hydrophobic and charged residues within the BH3 domain. In particular, the interaction of the hydrophobic H4 residue with BCL-xL shows a distinct behavior (L148 of PUMABH3 interacts with E96 and W137, while L148 of dNPUMA interacts with N136). This difference could be attributed to the TA of dNPUMA, which allowed Y152 and E158 to interact with the BCL-xL globular domain (Figure 4). We hypothesize that the allosteric relationship between the TAs of BCL-xL and PUMA guides the positioning of the PUMA BH3 IDR within the BCL-xL groove, following an in-duced-fit mechanism as proposed by Bekker *et al*. (2023). This results in a BH3-in-groove conformation distinct from that obtained with the BH3 domain alone, from which the BH3m molecules were designed. Consequently, the allostery highlighted in our study leads to a conformation where competitive inhibition by BH3m is less effective. Our detailed mapping re-examines the relative importance of specific residues within the BH3-in-groove of doubly membrane-anchored PUMA / BCL-xL and suggests alternative potential sites for therapeutic targeting.

**Figure 7:**
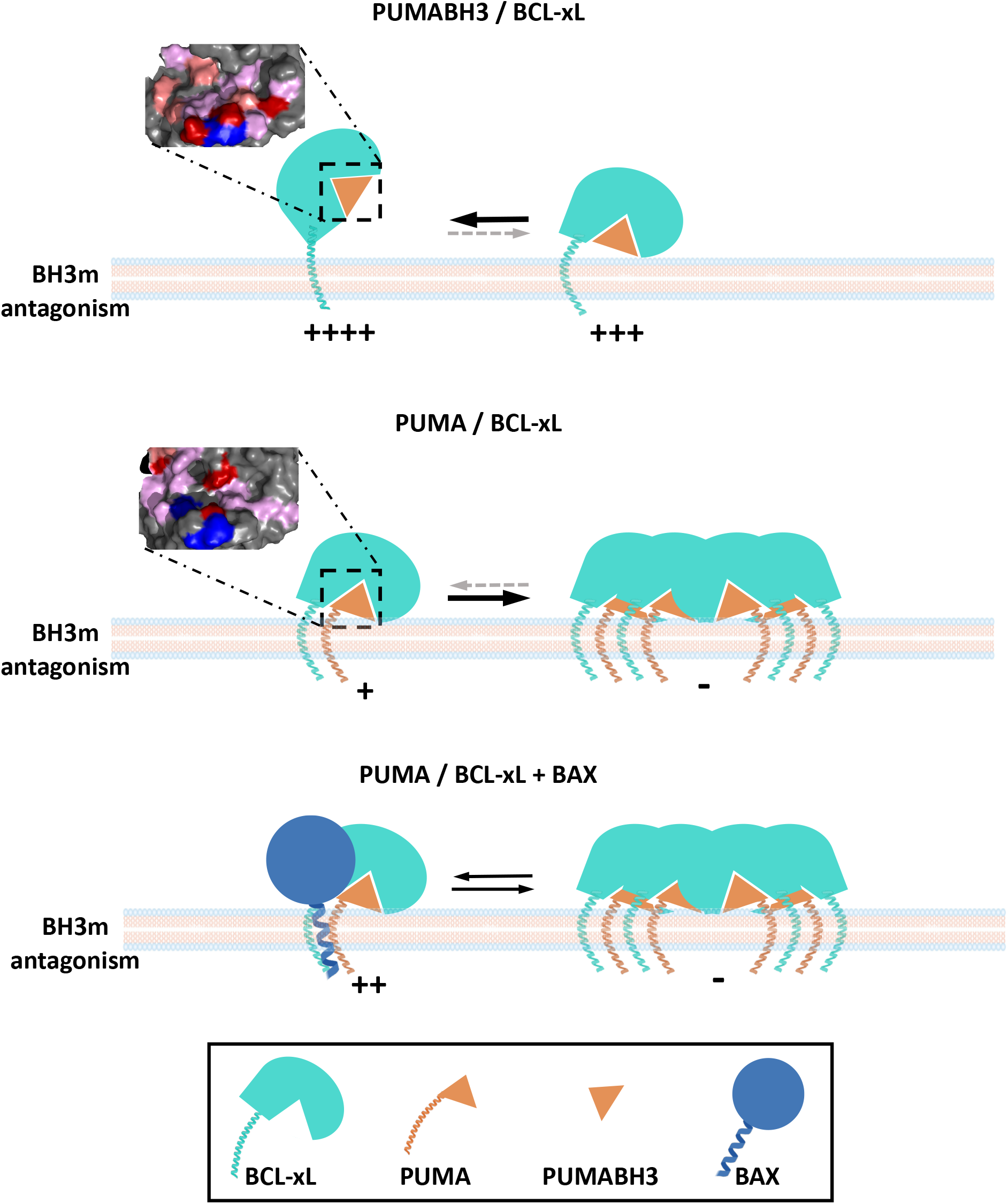
Schematic overview of allosteric regulation of PUMA / BCL-xL BH3-in-groove interface with their tail anchors The orientation of the BH3 binding interface towards the membrane is driven by the curvature of the BCL-xL TA within the membrane (top panel) and further stabilized by the presence of PUMA TA (middle panel), which maintains a double-kinked conformation of both TA helices. BCL-xL and PUMA TA allosterically modify the BH3 binding interface (see the enlarged interfaces in the top and middle panels) and also favors the formation of high molecular weight protein complexes, in a situation where BH3m is less competitive for binding antagonism. Potential mitochondrial proteins associated with PUMA/BCL-xL high molecular weight complexes are not shown. In addition, PUMA binding to BCL-xL increases the proximity of BAX to BCL-xL complexes (bottom panel), which may limit high molecular weight assembly. BAX promotes the sensitivity of PUMA / BCL-xL to BH3m antagonism through its own TA and its ability to interact with BCL-xL TA. Note that the conformational consequences of this functional relationship remain to be determined.

Our biochemical analysis indicates that binding to membrane-anchored partners enriches BCL-xL at subcellular membranes, consistent with the notion that engagement of BCL-xL groove by protein ligands enhances its steady-state membrane anchoring (Bleicken *et al*, 2017 and references therein) and reduces its dynamic shuttling between subcellular membrane and the cytosol (King *et al*, 2022). This stabilization in membrane likely contributes to maintain complexes in a BH3m resistant configuration. Displacement from the groove of BCL-xL TA by the PUMABH3 domains, making the TA available for insertion into the membrane (Bhat *et al*, 2012; Yao *et al*, 2015), cannot solely account for changes observed here, as only PUMA (not PUMAdC) alters BCL-xL membrane localization. Our biochemical investigation of cells co-expressing PUMA and BCL-xL unravels new features of the BCL-2 interaction network in whole cells. Mass spectrometric IP analysis and BN-PAGE reveal indeed that the interactome of BCL-xL is richer and organized with greater complexity when it is engaged by membrane-anchored ligands. The apparent high molecular weights of PUMA / BCL-xL complexes indicate that our current view of initiators / BCL-xL complexes as heterodimers needs to be re-appraised to take into account that additional proteins are in the proximity of PUMA / BCL-xL in whole cells, in agreement with previous reports (Singh *et al*, 2017; Bogner *et al*, 2020). Higher-order BCL-xL complexes have been observed to be mediated by the TA of BCL-xL (Bleicken et al, 2017). Doubly membrane-anchored complexes might allow currently underestimated binding interfaces in PUMA or BCL-xL to be manifest, possibly favored by the conformational constrains described above. We note that our mass spectrometric analysis identified, in BCL-xL pull downs in the presence of PUMA, TOM40 (the core component of the mitochondrial outer membrane translocation machinery) which was described to interplay with BCL-2 family members (Lalier *et al*, 2022) and several components of the mitochondrial inner membrane, including members of the cytochrome C oxidase complex (Figure S5D). Reports of interactions between PUMA and the mitochondrial inner membrane pyruvate carrier (Kim *et al*, 2019), of interactions between BCL-xL and the F1F0 ATP Synthase (Alavian *et al*, 2011) or even cytochrome C (Kharbanda *et al*, 1997) have already put forth contacts between BCL-2 family members and proteins of the mitochondrial intermembrane space or inner membrane. One may speculate from our data that doubly membrane-anchoring favor such interactions, leading to metabolic adaptation in live stressed cells (Green, 2019).

Notably, the effects of membrane-anchored PUMA binding might not be restricted to BCL-xL as it also promotes the mitochondrial assembly of endogenous MCL-1 into high molecular weight protein complexes. Ectopic PUMA distributes between an excess of BCL-xL and endogenous MCL-1 (Figure S5F) and, even though we cannot formally rule out that fractions of MCL-1 and BCL-xL may be proximal upon expression of PUMA, the absence of detectable MCL-1 in BCL-xL pull downs suggests that these two proteins assemble with PUMA essentially in distinct multimolecular complexes. Moreover, in agreement with our results, FRET signals associated to PUMA / BCL-xL and PUMA / MCL-1 interactions in whole cells resisted to BH3m treatments (Osterlund *et al*, 2022). Thus, the synergistic effects of dual inhibition of BCL-xL and MCL-1 by BH3m, efficient across a wide range of cell lines, may nevertheless be significantly limited when membrane anchoring of pro-MOMP ligands prevent displacement.

The observation that co-expression of PUMA and BCL-xL enhances BAX (and BAK) recruitment close to BCL-xL contradicts the widely accepted idea that these executioner proteins compete with initiators for binding to BCL-xL. Instead, it argues that the promotion of doubly anchored complexes shifts the BCL-xL network towards a new equilibrium state leading to increased proximity of other pro-MOMP proteins of the BCL-2 family, due to the involvement of specific regulatory motifs. Engagement with PUMABH3 preferentially occupies the BCL-xL groove over to that of BAX or BAK, based on the affinities of their respective BH3 domains (Chen *et al*, 2005). We assume that the formation of PUMA / BCL-xL complexes, while stabilizing BCL-xL at subcellular membranes, prevents the retrotranslocation of mitochondrial BAX into the cytosol. This process indeed requires canonical BAX / BCL-xL binding at the membrane (Edlich et al, 2011). Consequently, upon PUMA co-expression with BCL-xL, BAX (and BAK) molecules tend to accumulate in the proximity of mitochondrial BCL-xL, due to aborted retrotranslocation, promoting the occurrence of non-canonical contacts. In return, BAX and BAK may interfere with the formation of the high molecular weight complexes observed upon co-expression of membrane-anchored PUMA and BCL-xL, as suggested by our BN-PAGE analysis of PUMA / BCL-xL complexes in BAX/BAK knock out cells (Figure 5D).

Dimerization and heterodimerization between the TAs of BAX and the guardians proteins represent an additional layer of regulation to the BCL-2 network of interactions and, specifically, the TAs of BCL-xL and BAX were shown to interact with each other in membranes (Andreu-Fernández *et al*, 2017; García-Murria *et al*, 2020; Lucendo *et al*, 2020; Beigl *et al*, 2024; Nguyen *et al*, 2024). A strong body of evidence advocate for the involvement of such an interplay, rather than a competition for in groove binding, in the BAX mediated regulation of PUMA / BCL-xL sensitivity to BH3m. Firstly we detected a proximity between the sole TA of BAX and BCL-xL in PUMA / BCL-xL expressing cells, indicating that this interaction is not antagonized by engagement of PUMA by BCL-xL. Secondly, this domain sensitizes cells expressing doubly anchored PUMA / BCL-xL complexes, but not PUMAdC / BCL-xL complexes to BH3m induced cell death. Thirdly swapping BCL-xL TA with that of TOM 20 (which does not interact with TAs of BCL-2 family members) or with that of BAX (which self-associates) dampens and enforces, respectively, the effects of endogenous BAX on the BH3m sensitivity of PUMA / BCL-xL complexes. We propose that BAX TA physically interferes with the adjustment of BCL-xL TA in the membrane, which, as inferred from dynamic modeling, is required for allosteric regulation of the BH3 binding site. Such regulation would favor BH3m antagonism of doubly anchored complexes and impede the formation of higher molecular complexes (Figure 7). Interestingly, missense mutation (G179E) in a residue of BAX crucial for its dimerization with BCL-2 homologs TAs promotes resistance to ABT-199 BH3m (Fresquet *et al*, 2014), suggesting a similar regulation for the formation of BCL-2 complexes. We assume that the relative limited impact of endogenous BAK (compared to that of BAX) stems from the weaker capacity of its C terminal end to engage with that of BCL-xL (Duart *et al*, 2023).

The influence of BAX on BH3m sensitivity of PUMA / BCL-xL complexes highlight an unexpected feedback effect of this executioner on the whole network of BCL-xL interactions. It defines BAX as a major antagonist of BCL-xL, capable of acting at different steps (akin to a dominant negative form of BCL-xL) to ensure complete mitochondrial permeabilization. On this aspect, our study updates the « rheostat » model » which postulated in 1993 (Korsmeyer *et al*, 1993) that death/survival decisions are critically determined by the antagonistic activities of multi-domain executioners versus pro-survival BCL-2 homologs. Our observations imply that transcriptional regulation of executioner expression might significantly impact on the predisposition to MOMP induced cell death. Examples include c-Myc driven BAX/BAK expression in young tissues, which was shown to link developmental growth to cell death (Sarosiek *et al*, 2017) or p53-regulated BAX expression in multiple myeloma samples, which was shown to correlate to BH3m induced cell death (Durand *et al*, 2024). Since BAX regulation of BH3m effects seems to require a localization of BAX at subcellular membranes (in order to interfere with the allosteric effects of TAs), one may infer from our studies that low noise BAX activation might be associate with BCL-xL dependency and favor BH3m efficiency. Consistent with this notion, an investigation using the Dependency Map portal reveals that CRISPR invalidation of BAX has a robust inverse correlation with the impact of CRISPR invalidation of BCL-xL on the fitness of various cell lines from diverse origins, and tends to positively correlate with sensitivity to WEHI-539, which itself is correlated with the effects of BCL-xL invalidation (Figure S7).

Our findings should prompt a reevaluation of our understanding of BCL-2 family proteins based on information derived from experimental models in soluble conditions or in which the proteins lack TA or where cellular background lacks BAX/BAK expression. They advocate for experimental strategies that should consider the intricate conformational dynamics influenced by the TA within lipid membranes and for approaches that consider the entirety of the interaction network, rather than focusing solely on isolated interactions. Our study opens up a number of avenues for targeting the network of interactions engaged by BCL-xL to promote cell survival. Firstly, it encourages the design of small molecule antagonists of BCL-xL based on the BH3-in-groove binding interface configuration of BCL-xL adopted following the allosteric regulation by TA reported here: such molecules may offer selective antagonism of BCL-xL complexes (e.g. involving PUMA), potentially avoiding adverse effects due to the disruption of other complexes (i.e. BAK/BCL-xL in platelets, Josefsson *et al*, 2020). Secondly, it provides a strong rationale for the recently reported synergy between BAX activating molecules, leading to its integration into the membrane, and BCL-xL antagonists (Lopez *et al*, 2022): this combination would be highly synergistic not only because BAX is an executioner of MOMP, but because it amplifies the biochemical events triggered by BCL-xL antagonism upstream of full activation. Thirdly, it implies that chemical or peptide mimics of interactions between BCL-xL, PUMA and/or BAX TAs should modulate BH3m efficiency. In particular, interfering with this TAs-mediated allosteric regulation could modulate the BH3-in-groove conformational equilibrium and improve protein druggability, as recently proposed (Mizukoshi *et al*, 2020). It is particularly relevant to note here that disrupting BCL-xL TAs dimerization was recently shown to reduce its pro-survival activity (Duart *et al*, 2023). Such line of research has the potential to uncover novel approaches to modulate the role of BCL-xL in the regulation of cell survival.

### Limitations of the study

Here we demonstrate allosteric regulation mediated by the TAs at the BH3-in-groove binding of BCL-xL and PUMA. While BRET technology avoids the need for biochemical approaches to evaluate BCL-xL antagonism in a cellular context, particularly membrane localization, it has its own limitations. The overexpression of fusion proteins with RLuc or YFP required for BRET experiments, although the functionality of these fusion proteins has been demonstrated elsewhere (Liu *et al*, 2019), could affect protein stability, intracellular localization, or interactions due to altered protein concentrations within cells. The use of genome editing to tag endogenous BCL-xL is certainly one way to address this issue.

Our molecular dynamics simulations are not entirely based on experimental structural biology data, particularly regarding the insertion of TA into membranes. Furthermore, the structure of BCL-xL does not account for the intrinsically disordered region in its N-terminal region. The unstructured N-terminal segment preceding the BH3 domain of PUMA was also intentionally excluded from this investigation as its contribution to BH3m antagonism is negligible according to our study. Despite these constraints imposed by our initial model and the fact that our molecular dynamics simulations encompass a finite time frame to 2 µs, we have uncovered significant differences that arise when considering the presence or absence of PUMA’s TA. Further structural studies, such as cryo-microscopic approaches, could be helpful in refining our conclusions.

We also wish to point out that we modeled a membrane with a fixed, standard lipidic composition. Our whole approach can nevertheless be used to study how changes in lipids influence the conformational dynamic of BCL-xL, as reported in (Tyagi *et al*, 2021). Further evaluations will allow to determine the influence of mixed lipid composition.

## MATERIALS AND METHODS

### KEY RESOURCES TABLE

List of source and identifier for reagents and resources used in this study. Links to all uploaded data are listed.

## RESOURCE AVAILABILITY

### Corresponding author contact

Further information and requests for resources and reagent should be directed to and will be fulfilled by the corresponding author Laurent Maillet.

## Material availability

Generated plasmids in this study are available on request to the corresponding author.

## EXPERIMENTAL MODEL

### Cell culture

All cell lines (MCF-7, HCT116 wild type and derivatives) were cultured at 37°C and 5% CO2 in RPMI 10% fetal bovine serum (FBS) + 1% penicillin-streptomycin and split 1 in 10 every 3 to 4 days. Cells were subjected to regular mycoplasma testing. BAX^CRISPR-/-^ and BAK^CRISPR-/-^ denote the inactivation of endogenous genes using virally delivered CRISPR-Cas9 technology. Lentivirus were produced from HEK 293T cells transfected with the pLentiCRISPRV2-Puro plasmid containing the guide sequences AGTAGAAAAGGGCGACAACC and GCCATGCTGGTAGACGTGTA, respectively. Cells were selected using 1 μg/ml puromycin and gene invalidation were confirmed by Western blot for the absence of detectable BAX and/or BAK compared the parental cells. _G_BCL-xL, _F_PUMA / _G_BCL-xL, _F_PUMAdC / _G_BCL-xL denote stable cell lines obtained after infection with lentivirus produced from LZRS plasmid encoding GFP-BCL-xL, FLAG-PUMA-2A-GFP-BCL-xL or FLAG-PUMAdC-2A-GFP-BCL-xL, respectively. Infected cells were selected with 1 mg.ml^-1^zeocin for 2 weeks and then sorted by Facs ARIA III to ensure homogenous GFP expression. _MB_CBAX denote stable cell lines obtained after infection with lentivirus produced from pLVX-EF1α-IRES-Puro-Myc-BirA-CBAX plasmid encoding Myc-BirA-CBAX (BAX last 23 amino acids) and puromycin selection (1 μg/ml). The expression of each transgene was verified by Western blot. For transient expression, cells were transfected at 70 to 80% confluence using Opti-MEM reduced serum medium with Lipofectamine 2000 according to the recommendations of the supplier.

### DNA constructs

The plasmids used in this study are listed in the key resources table. Cloning details will be provided upon request. All of the proteins are expressed from human cDNA and, for tBID, also from mouse cDNA. Plasmids for transient expression of untagged protein or domain were cloned into the pcDNA3 backbone, RLuc and YFP fused to the BCL-2 family member were assembled into the peYFP-C1 and pRLuc-C1 backbones, respectively. Substituting YFP with mCherry or GFP in the pEYFPC1 plasmid resulted in mCherry or GFP-fused proteins, respectively. In the plasmid for full length or truncated PUMA expression, we exchanged its BH3 binding domain with the BH3 sequence of BID (Supplementary Table 1) using in-Fusion HD cloning. Site-directed mutagenesis was used to introduce mutations described in the study. All DNA manipulations and plasmids were confirmed by sequencing before use. To generate stable cell lines, transgenes were cloned into LZRS retroviral or pLVX-EF1α-IRES-Puro lentiviral backbones.

The nomenclature used throughout this study indicates proteins fused with R-Luciferase by a subscripted R, with YFP by a subscripted Y, with GFP by a subscripted G, with Flag by a subscripted F, with mCherry by the subscripted mC, and with MycBirA by the subscripted MB.

## EXPERIMENTAL PROCEDURES

### BRET saturation curve and dose-response

BRET experiments were performed as described in (Pécot *et al*, 2016). Briefly, cells were plated in 24-well plates and transfected with increasing amounts (25–500 ng/well) of plasmids coding for a BRET acceptor (protein fused to YFP), and constant amounts (25 ng/well) of plasmid expressing a BRET donor (protein fused to R-Luciferase). BRET measurement was performed using the lumino/fluorometer Mithras LB 940 (or LB 943 model) Berthold Technologies, France) after washing once with PBS and injection of coelenterazine H substrate in wells in PBS at a final molarity of 5 μM (Interchim). BRET signal corresponds to the emission signal values (530 nm) divided by the emission signal values (485 nm). The BRET ratio was calculated by subtracting the BRET signal value obtained with co-expressed donor and acceptor by that obtained with the donor protein co-expressed with untagged donor protein. Data shown are representative of at least two independent experiments for saturation curves. For dose-response analysis, transfected cells were harvested and seeded in triplicate in 96-well white plates. Twenty-four hours later, cells were treated with a concentration range 1:2 serial dilution of BH3m (0,0006 to 10 μM) for 16 hours, and BRET was measured as described above. Data are presented as the results of at least three independent experiments.

### Imaging

Cells cultured on the glass side were transfected with mCherry-fused PUMA, and then loaded with MitoTracker Deep Red for 30 minutes in culture medium at 37°C. They were then washed twice with PBS, fixed in PBS containing 4% paraformaldehyde/4% sucrose for 15 minutes and mounted with ProLong Diamond Antifade Reagent. Fluorescence images were acquired with Nikon A1 Rsi Inverted Confocal Microscope (Nikon, Tokyo, Japan) with NIS-Elements software (Nikon).

### Modelling of BCL-xL structure in complex with PUMABH3 or dNPUMA

Membrane-anchored BCL-xL was obtained by merging the known NMR BCL-xL structure (PDB Id: 2LPC) with an ideal alpha helix model for BCL-xL *α*9 helix (residues S203 to K233)(Lee & Fairlie, 2019). PUMABH3 domain bound membrane-anchored BCL-xL was derived from the structure of globular BCL-xL in complex with PUMABH3 peptide (PDB Id: 2M04 from (Follis *et al*, 2013). In the NMR structure 2M04, PUMABH3 (residues E130 to R154) are already adjusted to BCL-xL, we added the tail anchor part of BCL-xL with an ideal alpha helix model for BCL-xL *α*9 helix (residues S203 to K233) to obtain BCL-xL. Finally, the doubly anchored dNPUMA / BCL-xL was modeled in four steps: (i) 2M04 was taken as a starting structure for the globular part, (ii) the two TA (BCL-xL: S203 to K233, PUMA: S166 to N193) relative orientation was studied by protein-protein docking methods (Pierce *et al*, 2011), (iii) BCL-xL was reconstituted in membrane context, which allowed to orient PUMA TA relative to PUMABH3, and (iv) and by merging to the remaining linker between PUMABH3 and PUMA TA (^155^RRQEEQQRHRP_165_). PUMA linker made of 11 amino acids showed no structure preference using standard secondary structure prediction or protein-protein analysis on BCL-xL, it was thus modeled as a coil structure. Note that this structure lacks the N-terminal (for which no structural data exist), which we found to have no direct effect on BH3m sensitivity. Each system was inserted in a POPC membrane.

At the beginning of the simulation both TA were tilted in the membrane (about 15° (Yao *et al*, 2015) and TA were parallel according to our modeling procedure. Charges and protonation states of the model were assessed using PROPKA (Olsson *et al*, 2011) at pH 7. Insertion in a POPC (Phosphatidylcholine) membrane was performed using packmol-memgen (Schott-Verdugo & Gohlke, 2019). The details of the system are given in Table 1.

**Table 1:** System preparation system sizes

| System | Initial structure | Atoms | Na+ | Lipids | Size (Å) | Simulation time (ns) |
| --- | --- | --- | --- | --- | --- | --- |
| PUMABH3 / BCL-xL | 2LPC/2M04 | 99 578 | 8 | 235 | 88.99 x 89.22 x 134.92 | 2000 |
| dNPUMA / BCL-xL | 2LPC/2M04 | 100 249 | 8 | 235 | 87.23 x 87.29 x 132.97 | 2000 |

### Molecular dynamics simulations

Molecular dynamics simulations were performed using AMBER 20 (Case *et al*, 2023), generating trajectories spanning at least 2 microsecond using ffSB14 force field for proteins (Maier *et al*, 2015) and lipids17 force fields for lipids, TIP3P water model (Price & Brooks, 2004), employing the recommended settings as described in literature (Dickson *et al*, 2022). Briefly, systems were modeled under periodic conditions, employing Particle Mesh Ewald (PME) electrostatics calculations (Darden *et al*, 1993), ffSB14 force field with a 10 angstrom cutoff and Langevin thermostat with a thermostat coefficient frequency of 1.0 ps_-1_. Upon protein insertion into the membrane, the system underwent minimization, comprising 5000 steps of steepest descent followed by 5000 steps of conjugate gradient, with a constraint of 2 kcal.mol_–1_.Å_–2_ applied to both proteins and lipids atoms. System heating was performed in NVT conditions where a constraint of 10 kcal.mol_–1_.Å_–2_ was applied to the atomics positions of proteins and lipids during the transition from 0K to 100K over 50ps and with a constraint of 5 kcal.mol_–1_.Å_–2_ from 100K to 303K for 100ps. System volume was equilibrated in NVT conditions for 100 ps using a restraint of 5 kcal.mol_–1_.Å_–2_, followed by a further equilibration period of 500 ps in NPT conditions under the same restraints conditions. Unrestrained simulations were then computed unrestrained for 2 microseconds.

### Molecular dynamics analysis

All simulations were visualized using PyMOL (https://pymol.org) and VMD (Humphrey *et al*, 1996). Data analysis were performed using XmGrace (https://plasma-gate.weizmann.ac.il/Grace/), Python (van Rossum & de Boer, 1991) and matplotlib (Hunter, 2007).

### Detailed energetic contribution analysis

System trajectories were analyzed using the MM/GBSA single trajectory free energy calculations using MMPBSA.py (Miller *et al*, 2012). The 600-2000 ns interval of the trajectories was considered with a calculation every 10 frames using IGB 5 and the LCPO method for surface calculation. Per-residue energy decomposition was obtained by adding 1-4 interactions to electrostatics and Van Der Walls energies. Further analyses were performed using in-house python programs and plotted using matplotlib.

### Subcellular fractionation

The heavy membrane and cytosol fractions from MCF7 and HCT116 cells were prepared by subcellular fractionation based on differential centrifugation. Cells were scraped, centrifuged, and washed twice at 1,000g for 10 minutes at 4°C with ice-cold PBS. They were then resuspended in volume-per-volume cell extraction buffer (MB-EDTA) containing 210 mM mannitol, 70 mM sucrose, 10 mM HEPES (pH 7.5), 1mM EDTA (pH7,5), 1 mM PMSF, and 1 tablet of protease inhibitor. Cells were lysed using a Dounce homogenizer on ice, and sequential centrifugation (500g for 5 minutes, 2×2000g for 5 minutes, and 13,000g for 10 minutes) resulted in pelleting the crude heavy membrane fraction. The supernatant was then centrifuged at 20,000g for 30 minutes at 4°C, and the resulting supernatant is the supernatant fraction. The crude heavy membrane pellet was washed once in cold MB-1mM EDTA (pH7,5) buffer and centrifuged at 500g for 3 minutes at 4°C. The supernatant was centrifuged at 10,000 g for 10 minutes at 4°C, and the resulting pellet, containing mitochondria and endoplasmic reticulum, was solubilized in MB-EDTA buffer.

### Western blot and BN-PAGE analysis

Proteins were obtained by lyzing cells with RIPA buffer (5 mM Tris-HCl pH 7.6, 150 mM NaCl, 1% NP-40, 1% sodium deoxycholate, 0.1% SDS). Protein samples were subjected to Mini-PROTEAN® TGX™ Precast Protein Gels, 4–20% and transferred to nitrocellulose membranes using Trans-BlotR Turbo™ Transfer System Cell system (Bio-Rad). The membrane was then blocked in 5% nonfat dry milk TBS 0.05% Tween 20 and incubated with primary antibody overnight at 4°C (listed in the key resource table). Blots were incubated with the appropriate secondary antibodies for 1 h at room temperature and visualized using the FUSION - FX 7 (Vilber).

Analysis of native protein complexes in membranes was performed by blue native polyacrylamide gel electrophoresis (BN-PAGE) following the supplier’s instructions (Life Technologies). Briefly, heavy membrane fractions were solubilized in 1% digitonin NativePAGE sample buffer (50 mM BisTris, 6N HCl, 50 mM NaCl, pH 7.2, 10% glycerol, 0.001% Ponceau S) for 10 minutes on ice. Insoluble material was removed by centrifugation (20,000 x g, 20 minutes, 4°C). After the addition of NativePAGE G-250 Sample Additive, the heavy membrane extracts were applied to blue native PAGE Novex 4-16% Bis-Tris gels and subsequently blotted onto a PVDF membrane for Western blot analysis.

### Incucyte imager-based videomicroscopy analysis

Cell death analysis was determined using an Incucyte Zoom imaging system (Sartorius). Cells were plated in 96-well plates and then treated with a concentration range 1:4 serial dilution of A-1331852 (0 to 10 μM) with and without S-63845 (0,5 μM) and AnnexinV-CF®594 Conjugate dye were then added. Plates were imaged every 60 minutes during 13h. The quantitative analysis was done using Incucyte cell-by-cell analysis software module (Sartorius).

### Immunoprecipitation assay

Protein lysates were obtained by lysing cells with IP buffer (TRIS 20mM pH7.5, 150mM NaCl, 1mM EDTA, 1% CHAPS, 1mM PMSF supplemented with protease and phosphatase inhibitors) for 45 minutes on a roller in the cold room and clarified at 13,000g for 15 minutes at 4°C. Immunoprecipitation was performed on 500 µg protein lysates incubated with 5 µL anti-FLAG® M2 magnetic beads or ChromoTek GFP-trap® magnetic agarose for 45 minutes on a cold roller, followed by washing on a magnetic separator. Pulled-down proteins were analyzed by Western blot or mass spectrometry.

### Mass Spectrometry analysis

Beads containing protein complexes were re-suspended in 50 mM NH4HCO3, pH=8. 10 mM DTT were added for 15 minutes at 50°C for reduction and then 10 mM MMTS were added at room temperature for alkylation during 10 minutes. Trypsin was added at the protein/trypsin ratio of 50:1. Digestion was performed overnight at 37 °C. Peptides were acidified to a final concentration of 1% formic acid (FA), followed by desalting using C18 StageTips. Micro BCA assay was performed and peptides were vacuum-dried in a SpeedVac concentrator and stored at -20°C until measured by LC-MS/MS. Immediately prior to micro-LC, the fractions were resuspended in H2O with 0.1% v/v formic acid at a concentration of 1 µg µL−1.

Each sample (5 µg) was separated into a micro 2D-LC 425 system (Eksigent) using a ChromXP C18CL column (3 μm, 120 A, 15 x 0.3 cm, Sciex) at a flow rate of 5 μL/minute. Water and ACN, both containing 0.1% formic acid, were used as solvents A and B, respectively. The following gradient of solvent B was used: 0 to 5 minutes 5% B, 5 to 75 minutes 5% to 35% B, then 10 minutes at 95% B, and finally 10 minutes at 5% B for column equilibration. As the peptides eluted, they were directly injected into a hybrid quadrupole-TOF mass spectrometer Triple TOF 5600+ (Sciex, Redwood City, CA, USA) operated with a ‘top 30’ data-dependent acquisition (DDA) system using positive ion mode. The acquisition mode consisted of a 250-ms survey MS scan from 400 to 1250 m/z, followed by an MS/MS scan from 230 to 1500 m/z (75 ms acquisition time, 350 mDa mass tolerance, rolling collision energy) of the top 30 precursor ions from the survey scan. For the relative quantification by SWATH-MS acquisition, each sample (5 µg) was analyzed using the LC-MS equipment and LC gradient described above for building the spectral library, but using a SWATH-MS acquisition method. The method consisted of repeating the whole gradient cycle, involving the acquisition of 32 TOF MS/MS scans of overlapping sequential precursor isolation windows (25 m/z isolation width, 1 m/z overlap, high-sensitivity mode) covering the 400 to 1200 m/z mass range, with a previous MS scan for each cycle. The accumulation time was 50 ms for the MS scan (from 400 to 1200 m/z) and 100 ms for the product ion scan (230 to 1500 m/z), giving a 3.5 s total cycle time. Peak extraction of the SWATH data was performed using SpectronautTM Pulsar X 13 software with default analysis settings. Retention time prediction type was set to dynamic iRT and calibration mode was set to automatic. These settings also included mutated decoy method and cross run normalization enabled. The FDR was estimated with the mProphet approach and Q-value cutoff for both precursor and protein was set to 1%13. Interference correction was enabled for quantification which required a minimum of 2 precursor-ions and 3 fragment-ions. The term protein refers to protein groups as determined by the algorithm implemented in SpectronautTM. The quantitative SWATH scores were processed by dividing the SWATH score by the GFP score in each sample to normalize the measurements against immunoprecipitated GFP-BCL-xL and to perform statistical comparative analyses of differentially interacting proteins using the t.test function (stats R package). Volcano Plots were generated using the corresponding R package, and ggplot2 was used to produce boxplot figures.

### Proximity Ligation Assay (PLA) flow cytometry experiment

_G_BCL-xL and _F_PUMA / GBCL-xL cells were collected by trypsinization and centrifugation, followed by fixation with 4% paraformaldehyde in PBS for 15 minutes at RT. Subsequently, cells were rinsed by PBS and permeabilized using 0.5% saponin in PBS for 25 minutes at RT. PLA staining was performed according to the manufacturer’s instructions (Sigma Aldrich #DUO92002 #DUO92004 #DUO94004). The primary antibodies used were anti-BAX (Enzo #4F11, 1/500) and anti-GFP (Abcam #ab290, 1/2000). In negative control cells, primary antibodies were omitted. Flow cytometry analysis was conducted on an Attune Nxt instrument (Thermo Fischer Scientific) in BL1 and RL1 channels for _G_BCL-xL expression and PLA signal, respectively. Data acquisitions and analyses were performed using the Attune Cytometric Software and statistical analyses were performed by Prism software.

### Flow cytometry analysis

Cytochrome-c release in MCF-7 cells was evaluated using the recombinant FITC-conjugated cytochrome C antibody (560263, BD Biosciences) after fixation/permeabilization using the FOXP3 Fix/Perm Buffer (Ebioscience, Thermofisher scientific). Flow cytometry analysis was performed on FACS Accuri C6 plus (BD Biosciences).

### BirA-CBAX interaction partner identification

_F_PUMA / _G_BCL-xL _MB_CBAX and _F_PUMAdC / _G_BCL-xL _MB_CBAX HCT116 express C terminal sequence BAX fused to Myc tagged BirA. Cells were growth in culture medium supplemented with 50 mM Biotin for 24 hour, and protein lysates were obtained by lysing cells with RIPA buffer. Total extracts were incubated with streptavidin agarose beads overnight at 4°C. Input and bead samples were resolved on a Mini-PROTEAN® TGX™ Precast Protein Gels, 4–20% and analyzed by Western blot for the indicated proteins.

## STASTISTICAL ANALYSIS

Statistical details of the experiments are provided in the respective figure legends. P < 0.05 was considered significant.

## ACKNOWLEDGEMENTS

We thank the members of the “Stress adaptation and tumor escape” laboratory for their support and helpful discussions. We thank Maximilien BERNE, Stéphane JEDELE and Anne-Emeline HUARD for setup and analysis of exploratory molecular dynamics simulations. We acknowledge the Cytocell - Flow Cytometry and FACS core facility (SFR Bonamy, BioCore, Inserm UMS 016, CNRS UAR 3556, Nantes, France) for their technical expertise and assistance, member of the Scientific Interest Group (GIS) Biogenouest and the Labex IGO program supported by the French National Research Agency (n°ANR-11-LABX-0016-01), and benefited from technical support from the IBISA MicroPICell facility (Biogenouest), member of the national infrastructure France-Bioimaging supported by the French National Research Agency (ANR-10-INBS-04). This research used resources of the GLiCID Computing Facility (Ligerien Group for Intensive Distributed Computing, https://doi.org/10.60487/glicid, Pays de la Loire, France). This work was supported by the INCa (SIRIC ILIAD INCa-DGOS INSERM-ITMO Cancer-18011 and PLBIO021-129NN), the departmental committee of LIGUE Contre le Cancer CD29 and CD44 and the Région des Pays de la Loire (“Trajectoire nationale” and PIRAMID project). P.P.J.’s laboratory is labeled by the Ligue Nationale Contre le Cancer (LNCC).

## AUTHOR CONTRIBUTIONS

Conceptualization: L.M., L.L., F.G., S.T. and P.P.J.; Supervision: L.M., S.T. and P.P.J.; Methodology: L.M., A.F., L.L., S.B.N., C.G. and S.T.; Validation: L.M., L.L., S.B.N., C.G., F.G. and S.T.; Formal analysis: L.M., A.F., L.L., S.B.N., C.G., F.G. and S.T.; Investigation: L.M., S.T. and P.P.J.; Writing: - Original draft L.M., S.T. and P.P.J. - Editing L.M. and P.P.J.; Visualization: L.M., L.L., F.G., S.T. and P.P.J.; Project administration: L.M. and P.P.J.; Funding acquisition: L.M. and P.P.J.

## DECLARATION OF INTERESTS

The authors declare no competing interest.

## Notes

### Competing Interest Statement

The authors have declared no competing interest.

